# Endosomal membrane state governs trafficking and cell migration under TFEB control

**DOI:** 10.64898/2026.09.02.748843

**Authors:** Julie Patat, Aline Mathey, Emna Ouni, Pallavi Mathur, Hugo Lachuer, Nadia Elkhatib, Mehdi Khaled, Gael Blivet-Bailly, Andrey S. Klymchenko, Julio Lopes Sampaio, Kristine Schauer

**Author notes:** Equal contribution (Julie Patat, Aline Mathey).

## Abstract

Transcription Factor EB (TFEB) regulates the biogenesis of lysosomes, which are acidic organelles of the endosomal network. While its role in cellular clearance is well established, the pleiotropic functions of TFEB across diverse cellular processes, as well as its activation in several cancer types, remain incompletely understood. Investigating TFEB in a bladder cancer model, we found that its depletion selectively impaired cell migration and adhesion. This phenotype was associated with the retention of the adhesion molecule integrin β5 (ITGβ5), in central intracellular compartments identified as multivesicular bodies (MVBs) upon TFEB knockdown (KD). Mechanistically, TFEB regulated cellular lipid composition and membrane fluidity of acidic endosomes that controlled ITGβ5 trafficking. Remarkably, exogenous supplementation with a monounsaturated fatty acid (MUFA) that increased MVB membrane fluidity was sufficient to phenocopy the intracellular trapping of ITGβ5 in 2D culture and 3D tumoroids. We further showed that TFEB-dependent maintenance of MVB membrane fluidity relies on ESCRT-0 subcomplex, HGS/Hrs, which retains cholesterol at MVBs. Together, these findings reveal how transcriptional programs shape endosomal membrane states to control cargo trafficking and cell behavior.

**Summary:** We uncover a previously unrecognized role for TFEB in controlling the lipid composition and membrane fluidity of multivesicular bodies, establishing endosomal membrane properties as a regulated output of transcriptional programs that govern cargo trafficking and cell behavior.

## INTRODUCTION

The endosomal network is traditionally viewed as a logistics platform that sorts signaling receptors, adhesion molecules, transporters, and their ligands between intracellular compartments and the plasma membrane, thereby shaping cell identity and behavior (Moreno-Layseca et al. 2019; Da Graça et al. 2025). Consistent with this central role, dysregulation of intracellular trafficking is implicated in a wide spectrum of human diseases, including cancer, neurodegenerative disorders, immune deficiencies, and metabolic diseases (Gilleron and Zeigerer 2023). Recent work reveals that endosomal sorting is not governed solely by canonical trafficking machineries (Gopaldass et al. 2024), but is instead dynamically tuned through crosstalk with cellular lipid homeostasis (Alonso-Bivou et al. 2025; Carpentier et al. 2025; Chen et al. 2025; Peng et al. 2025; Song et al. 2022) (*preprint* Otakhor et al. 2025). Although several molecular players at this interface have begun to emerge from these studies, a unifying conceptual framework explaining how membrane trafficking and lipid homeostasis are coordinated remains lacking.

One key regulator of the endosomal system is the transcription factor EB (TFEB), known as the master regulator of lysosomal biogenesis and autophagy (Settembre et al. 2011; Ballabio 2016). TFEB is a ubiquitously expressed member of the evolutionarily conserved microphthalmia/transcription factor E (MiT/TFE) family, which also includes MITF, TFE3, and TFEC (Hertwig 1942; Beckmann, Su, and Kadesch 1990; Carr and Sharp 1990; Zhao et al. 1993). The target genes of MiT/TFE factors comprise the CLEAR (Coordinated Lysosomal Expression and Regulation) gene network, which includes genes involved in lysosomal function and autophagy (Sardiello et al. 2009; Palmieri et al. 2011). TFEB shares considerable functional redundancy with TFE3, which similarly regulates late endosome and lysosome biogenesis and promotes autophagy. Both TFEB and TFE3 are activated in response to cellular stress signals such as nutrient deprivation (Martina et al. 2014), and multiple CLEAR network genes are co-regulated by these transcription factors (Huan et al. 2005, 2006; Settembre et al. 2013; Salma et al. 2015; Martina et al. 2016; Pastore et al. 2016).

In addition to its canonical role in lysosomal biogenesis and autophagy, recent studies have uncovered a broader regulatory function for TFEB in a context- and tissue-specific manner. These include the regulation of lipid metabolism in the liver (Pastore et al. 2017), as well as the induction of mitochondrial biogenesis and fatty acid oxidation through peroxisome proliferator-activated receptor alpha (PPARα) and its co-activator PGC1α in skeletal muscle (Erlich et al. 2018) and adipocytes (Evans et al. 2019). Additionally, TFEB has been implicated in the regulation of the Wnt/β-catenin signaling pathway, a key developmental and homeostatic regulator, both *in vitro* and *in vivo* (Calcagnì et al. 2016). These findings suggest that TFEB act as a convergent node, integrating environmental and metabolic cues to modulate a wide array of cellular processes.

TFEB has recently gained attention for its role in several cancers, including renal cell carcinomas (RCC) and pancreatic ductal adenocarcinoma (PDAC) (Perera et al. 2015; Perera, Di Malta, and Ballabio 2019; Wei, Testa, and Argani 2022; Zoncu and Perera 2023). In RCC, TFEB activation is driven by gene fusions or amplifications (Bakouny et al. 2022; Wei et al. 2022), and kidney-specific overexpression of TFEB in transgenic mice results in renal clear cell changes, severe cystic pathology, renal cysts, and papillary carcinomas with liver metastases (Calcagnì et al. 2016; Di Malta et al. 2023). In PDAC, nuclear localization of TFEB along with increased expression of TFEB target genes has been observed, yet in the absence of gene fusions (Perera et al. 2015; Zoncu and Perera 2023). We have previously shown that TFEB is also activated in aggressive bladder cancer cell lines (Mathur et al. 2023). Interestingly, while these aggressive cell lines exhibit a general enrichment of ‘CLEAR’ network, we did not observe the canonical TFEB-driven increase in lysosomal numbers compared to normal human urothelial cells. Instead, TFEB activation in this context primarily drives a dramatic shift in the subcellular positioning of acidic endosomes (Mathur et al. 2023). Given that bladder cancer affects over half a million people worldwide and incurs some of the highest treatment-related healthcare costs per patient, we sought to investigate the functional role of TFEB in this malignancy.

In this study, we uncover a previously unrecognized function of TFEB in regulating cellular membrane fluidity, a fundamental biophysical property that governs membrane organization, trafficking, and membrane protein function. We show that membrane fluidity of CD63-positive, acidic, late endosomes is increased upon TFEB depletion, which in turn controls the trafficking of integrin β5 (ITGβ5) and the migration of bladder cancer cells. We demonstrate that modulating endosomal membrane fluidity, either by supplementing cells with the unsaturated oleic acid or by inhibiting ESCRT-0 subunit HGS/Hrs, is sufficient to disrupt ITGβ5 trafficking. Together, we show that TFEB-dependent regulation of endosomal membrane fluidity functionally integrates cellular lipid metabolism with cancer migration through the trafficking of ITGβ5. These findings establish regulation of endosomal membrane properties as a critical and previously underappreciated layer of transcriptional control over cell behavior.

## Results

### TFEB regulates cell migration and trafficking of ITGβ5

Although constitutive TFEB activation is typically associated with enhanced lysosomal biogenesis, we previously demonstrated that TFEB activation in aggressive bladder cancer cells primarily drives anterograde lysosomal trafficking (Mathur et al. 2023), prompting us to investigate its noncanonical functions. We depleted TFEB by siRNA in the KU19-19 bladder cancer cell line, previously characterized by nuclear TFEB and a CLEAR gene signature (Mathur et al. 2023) and performed a RNASeq analysis on cells that showed 90% reduced TFEB levels (**SFig. 1A,B**). Gene Ontology enrichment analysis revealed that differentially expressed transcripts were primarily enriched for ribosomal proteins, together with gene sets associated with focal adhesions, respiratory chain complexes, and secretory and cytoplasmic vesicle compartments (**Fig. 1A**). Comparison with previously published TFEB-regulated gene lists in HeLa and HEK293T cells (Palmieri et al. 2011; Gambardella et al. 2020; Diamant et al. 2025) revealed that 84%, 16% and 7% of differentially regulated RNAs were also found in the Gambardella RNASeq, Palmery ChiPSeq, and the Diamant ChipSeq dataset, respectively. Our attention fell on the signature of ‘focal adhesions’ that was the most enriched one after ribosomal proteins in bladder cancer and additionally highly enriched in the Palmery ChiPSeq and Gambardella RNASeq that showed significant overlap with our dataset (**SFig. 1D, E**). We thus investigated cell migration in a wound healing assay using KU19-19 cells and found that TFEB KD led to about 50% reduction in the speed of the wound closure after scratching (**Fig. 1B,C**). We confirmed that cell proliferation was not induced by the wound scratching using the proliferation inhibitor mitomycin C (**SFig. 1F**). Additionally, cell spreading was reduced upon TFEB KD giving rise to a reduction in cell size (**Fig. 1D**).

**Figure 1.**
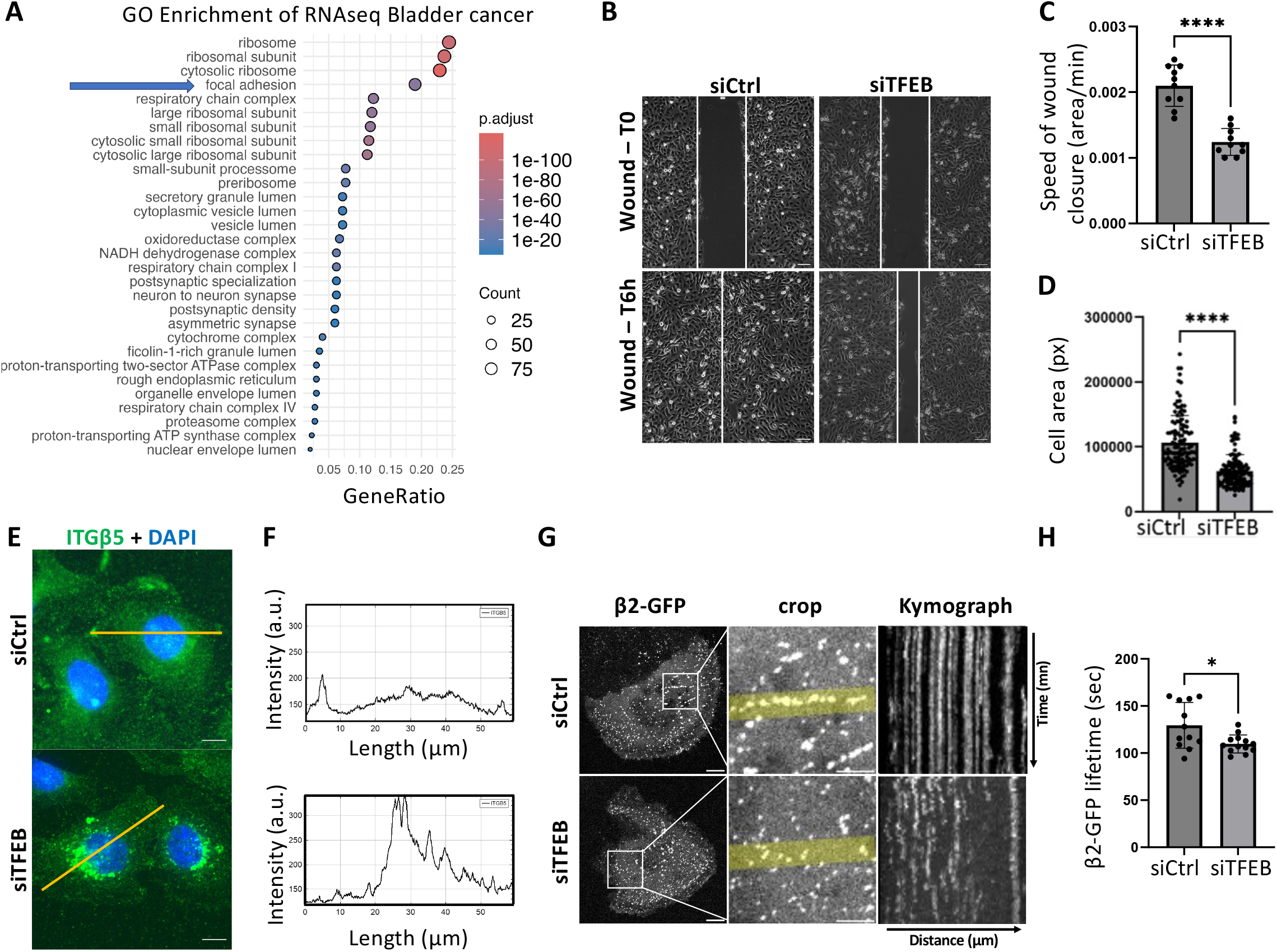
TFEB regulates cell migration and trafficking of ITGβ5. **(A)** Gene Ontology analysis of TFEB targets from RNA-seq data from KU19-19 cells. **(B)** Representative wound-healing assay of KU19-19 cells transfected with siControl (Ctrl) or siTFEB. Wound closure was monitored over 360 min (6 h). Scale bars, 100 μm. **(C)** Quantification of the speed of wound closure, calculated as |Tₙ–T₀|/T₀ in area/mins. Data represent mean ± SD from N=3; ****p < 0.0001 (Student’s t test). **(D)** Quantification of individual cell area in pixels for KU19-19 cells treated with siCtrl or siTFEB. Data represent mean ± SD from N=3; ****p < 0.0001 (Mann–Whitney test). **(E)** Immunofluorescence of ITGβ5 in KU19-19 cells after siCtrl or siTFEB treatment. Nuclei were counterstained with DAPI. Scale bars, 10 μm. **(F)** Line-scan density profile of ITGβ5 intensity corresponding to panels in (E). **(G)** Live-cell imaging of KU19-19 cells transfected with β2-GFP and either siCtrl or siTFEB for 48 h. Cells were imaged every 5 s for 5 min. Kymograph analysis of β2-GFP dynamics was performed using the KymographBuilder plugin in FIJI. Scale bars, 10 μm (overview) and 5 μm (kymograph). **(H)** Quantification of β2-GFP lifetime in seconds per cell. Data represent mean ± SD from N=3; *p < 0.05 (Student’s t test).

Integrins are major players of adhesion and migration, thus we next tested if addition of the integrin β3 and β5-specific RGD peptide, cilengitide (CLG), impacted migration. Whereas CLG inhibited migration speed in control KU19-19 cells, it did not further decrease speed in TFEB KD cells (**SFig. 1G**), indicating that integrin β3- and β5-dependent migration act under a common pathway with TFEB. Immuno blot analysis of several integrin proteins (ITBβ1, ITGβ3 and ITGβ5) in KU19-19 cell lysates showed no changes upon TFEB KD (**SFig. 1H**). Yet, visualization of the cellular localization of ITGβ5 by immunofluorescence revealed that it accumulated in central compartments of KU19-19 TFEB KD cells (**Fig. 1E,F).** ITGβ5 is known to inhibit dynamics of clathrin-coated pits due to the stabilization of flat clathrin lattice at the plasma membrane, also called clathrin-coated plaques (Baschieri et al. 2018; Elkhatib et al. 2017). Thus, we assessed the dynamics of the clathrin-adaptor β2-GFP. Pit lifetime was decreased (**Fig. 1G,H**) and β2-GFP speed dynamics were increased (**SFig. 1I,J**) upon TFEB KD, which indicated a loss of ITGβ5 at adhesive clathrin-coated plaques. Analysis of TFEB KD in JMSU1 cells (**SFig. 1A,C**), another bladder cancer cell line with nuclear TFEB and a CLEAR gene signature (Mathur et al. 2023), confirmed reduced migration under siTFEB (**SFig. 1K,L**). These results indicated that TFEB depletion impairs bladder cancer cell migration through intracellular sequestration of integrins.

### ITGβ5 is trapped in acidic endosomes marked by CD63

We have previously found that depletion of TFEB resulted in a perinuclear clustering of acidic LAMP1-positive endosomes in bladder cancer cells (Mathur et al. 2023). Our analysis with additional endocytic marker proteins in KU19-19 cells revealed that this compartment highly colocalized with CD63, a marker for multivesicular bodies (MVBs) that are acidic, late endosomes (**SFig. 2A,B**). Co-immunostaining of CD63 and ITGβ5 revealed a significant increase of their co-localization upon TFEB depletion (**Fig. 2A,B**), indicating that ITGβ5 accumulated in this acidic compartment. Whereas the CD63/LAMP1 positive endosomes were perinuclearly clustered upon TFEB depletion (**Fig. 2A,C**), the distribution of early endosomes (marked by EEA1) did not change (**SFig. 1C**). Perinuclear clustering of the CD63-positive endosomes upon TFEB KD was also observed in JMSU1 bladder cancer cells (**SFig. 2D,E**). Yet, total protein levels of the endosomal markers EEA1, CD63 and LAMP1 were unchanged under these conditions in both cell lines (**SFig. 2F**). In contrast, silencing of TFEB reduced the expression of the ESCRT-0 subunit HGS/Hrs in KU19-19 and JMSU1 cells (**Fig. 2D,E** and **SFig. 2G-I**), whereas other ESCRT components, including the ESCRT-I protein TSG101 and the ESCRT-III–associated protein ALIX, were not affected in both cell lines (**SFig. 2F**). Co-immunostaining of HGS/Hrs and CD63 upon siTFEB showed unchanged co-localization (**SFig. 2J,K**), but overall HGS/Hrs signal intensity was decreased (**SFig. 2L**). STED imaging of CD63 and HGS/Hrs showed that TFEB KD resulted in an enlargement of MVB in bladder cancer cells (**Fig. 2F, SFig. 2M**). Because HGS/Hrs plays a role in the formation of intraluminal vesicles (ILVs) of MVBs that give rise to extracellular exosomes (Edgar et al. 2014), we additionally assessed exosome abundance in cell culture supernatants of KU19-19 cells after enrichment and filtering. TFEB silencing markedly reduced exosome numbers by half (**Fig. 2G**), which was accompanied by a corresponding decrease in CD63 levels in cell culture supernatant detected by immunoblotting (**Fig. 2H**). Collectively, our data established TFEB as a key regulator of MVB homeostasis through HGS/Hrs, thereby governing ITGβ5 trafficking.

**Figure 2.**
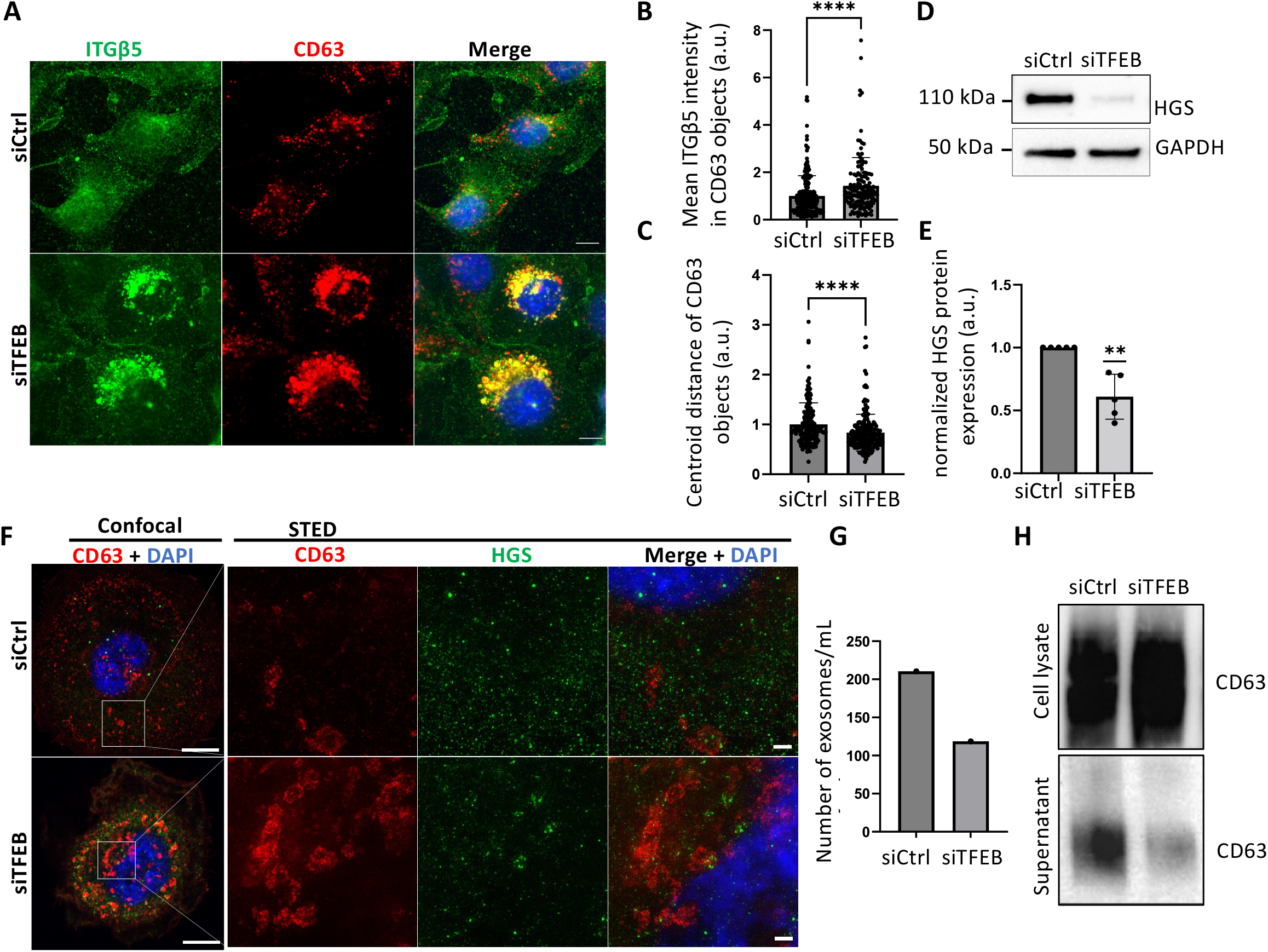
TFEB depletion promotes ITGβ5 accumulation in CD63-positive compartments and reduces HGS expression. **(A)** Immunofluorescence of ITGβ5 and CD63 (marker of multivesicular bodies, MVBs) in KU19-19 cells treated with siControl (Ctrl) or siTFEB. Nuclei were counterstained with DAPI. Scale bars, 10 μm. **(B)** Quantification of mean ITGβ5 fluorescence intensity within CD63-positive structures (arbitrary units, a.u.). Data represent mean ± SD from N=3; ****p < 0.0001 (Mann–Whitney test). **(C)** Quantification of the mean distance of CD63-positive structures from the cell centroid (arbitrary units, a.u.). Data represent mean ± SD from N=3; ****p < 0.0001 (Mann–Whitney test). **(D)** Immunoblot analysis of HGS in KU19-19 cells treated with siCtrl or siTFEB. GAPDH served as a loading control. **(E)** Densitometric quantification of HGS protein levels normalized to loading control (arbitrary units, a.u.). Data represent mean ± SD from N=4; **p < 0.01 (one-sample Student’s t test). **(F)** Images from stimulated emission depletion (STED) microscopy of CD63 and HGS/Hrs in KU19-19 cells treated with siCtrl or siTFEB. Nuclei were stained with DAPI. Scale bars, 10 μm (overview) and 1 μm (inset). **(G)** Quantification of the number of extracellular vesicles (exosomes) per ml in the supernatants of KU19-19 cells treated with siCtrl or siTFEB using nonaFCM. **(H)** Immunoblot analysis of CD63 in total cell lysates (upper panel) and supernatants (lower panel) from KU19-19 cells treated with siCtrl or siTFEB.

### ESCRT-0 HGS/Hrs regulates ITGβ5 trafficking downstream of TFEB

Next, we tested whether HGS/Hrs functions downstream of TFEB. HGS/Hrs was effectively depleted by siRNA in KU19-19 and JMSU1 cells after 48h (**SFig. 3A**) and resulted in slowed migration as well as reduced adhesion of both cell lines (**Fig. 3A-C; SFig. 3B-D**). Moreover, both cell lines showed perinuclear clustering of MVB upon HGS depletion, mimicking the siTFEB phenotype (**Fig. 3D,E; SFig. 3E,F**). Co-immunostaining of ITGβ5 and CD63 in KU19-19 cells revealed a significant increase of their co-localization upon HGS/Hrs depletion (**Fig. 3F**). Because anterograde trafficking of MVB towards the plasma membrane is specifically regulated by the small GTPase Rab27 (Ostrowski et al. 2010), we investigated RAB27 protein levels upon HGS/Hrs depletion. We found that RAB27 was reduced upon HGS/Hrs KD in both cell lines (**Fig. 3G,H; SFig. 3G,H**). To determine the functional relationship between TFEB and HGS/Hrs, we performed rescue experiments by transfecting siCtrl- and siTFEB-treated cells with HGS-RFP for 24 h (**Fig. 3I**). Overexpression of HGS/Hrs restored the peripheral distribution of CD63-positive endosomes in TFEB-depleted cells, promoting their relocalization toward the plasma membrane. These results indicate that HGS/Hrs can rescue the endosomal positioning defects caused by TFEB depletion and are consistent with HGS/Hrs functioning downstream of TFEB. Lastly, bioinformatic analysis of an aggressive bladder cancer patient cohort from the PRISM study (Pradat et al., 2023) performed at Gustave Roussy Hospital indicated a significant enrichment of the TFEB-dependent CLEAR signature in samples classified as ’high HGS’ compared to ‘low HGS’ (**Fig. 3J**). Indeed, we found a significant positive correlation between TFEB signature and HGS expression (**Fig. 3K**), indicating overlapping pathway involvement of both proteins. Together, these results indicated that HGS/Hrs regulates anterograde trafficking of MVBs and ITGβ5 via Rab27 downstream of TFEB.

**Figure 3.**
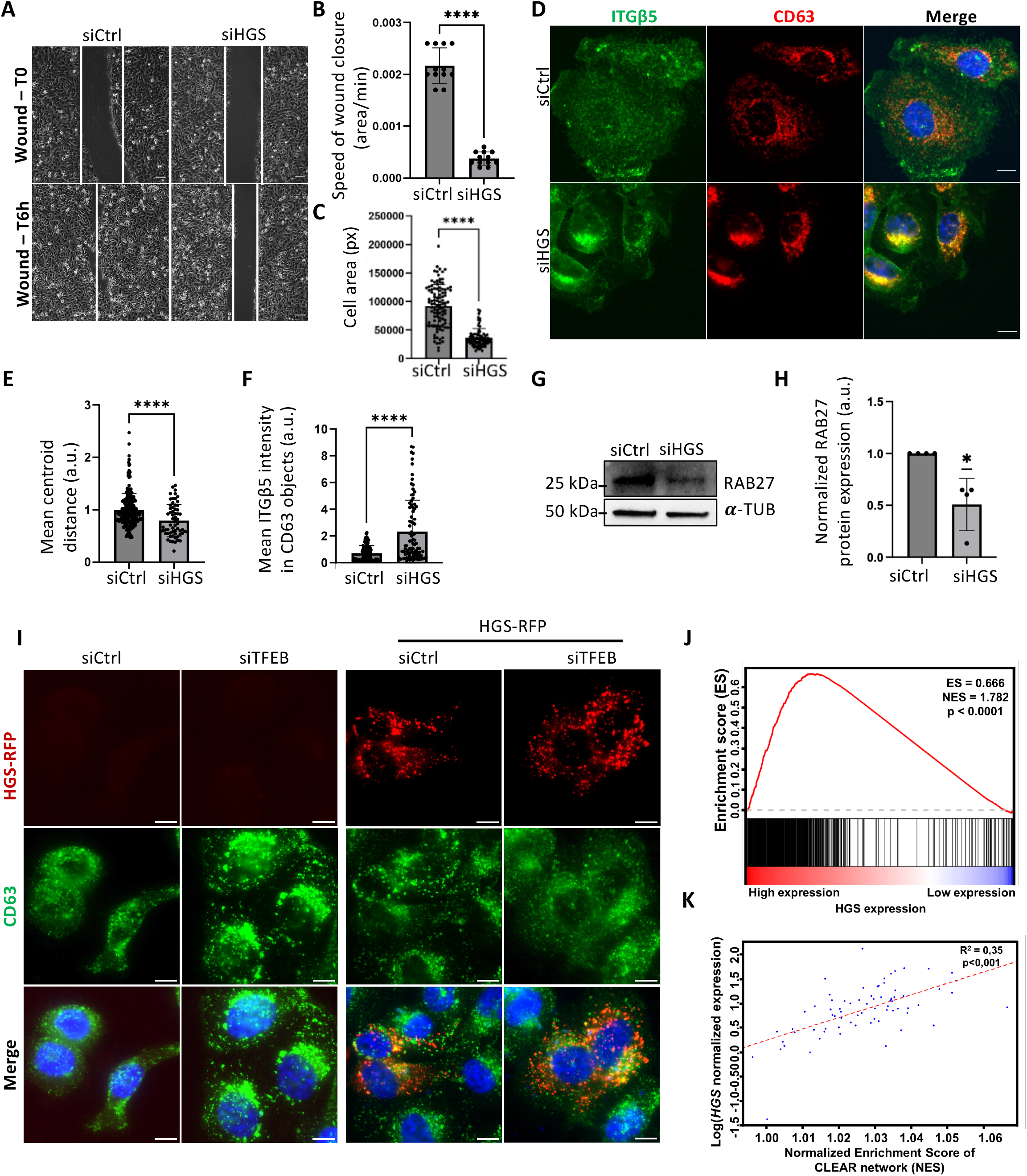
ESCRT-0 HGS/Hrs regulates ITGβ5 trafficking downstream of TFEB. (A) Representative wound-healing assay of KU19-19 cells transfected with siControl (Ctrl) or siHGS. Wound closure was monitored over 360 min (6 h). Scale bars, 100 μm. (B) Quantification of wound closure, calculated as |Tₙ–T₀|/T₀ in area/min. Data represent mean ± SD of N=3; ****p < 0.001 (Student’s t test). (C) Quantification of individual cell area in pixels (px) for KU19-19 cells treated with siCtrl or siHGS. Data represent mean ± SD of N=3; ****p < 0.0001 (Mann–Whitney test). (D) Immunofluorescence of ITGβ5 and CD63 in KU19-19 cells after siCtrl or siTFEB treatment. Nuclei were counterstained with DAPI. Scale bars, 10 μm. (E) Quantification of the mean distance of CD63-positive objects to the cell centroid, normalized to cell area, in KU19-19 cells treated with siCtrl or siHGS in arbitrary units (a.u.). Data represent mean ± SD of N=3; ****p < 0.0001 (Mann–Whitney test). (F) Quantification of mean ITGβ5 fluorescence intensity within CD63-positive objects, normalized to cell area in arbitrary units (a.u.). Data represent mean ± SD of N=3; ****p < 0.0001 (Mann–Whitney test). (G) Immunoblot analysis of Rab27 in KU19-19 cells treated with siCtrl or siHGS. α-Tubulin served as a loading control. (H) Densitometric quantification of RAB27 protein levels from immunoblots in KU19-19 cells treated with siCtrl or siHGS in arbitrary units (a.u.). Data represent mean ± SD of N=4; *p < 0.05 (One sample Student’s t test). (I) Immunofluorescence of CD63 in KU19-19 cells after siCtrl or siTFEB treatment and overexpression of HGS-RFP. Nuclei were counterstained with DAPI. Scale bars, 10 μm. (J) Gene Set Enrichment Analysis (GSEA) results showing the enrichment score (ES) of CLEAR network genes in bladder cancer biopsies with high vs. low HGS expression. The lower panels present the ranked Log2 Fold Change for each genes; red color indicates higher expression in the high HGS expression, while blue indicates lower expression in the low HGS expression group. Vertical black lines mark the positions of genes belonging to the CLEAR network. (K) Correlation between the logarithms of HGS normalized expression and the Normalized Enrichment Scores (NES) for the CLEAR network. NES have been computed individually for each bladder cancer biopsy and based on genes expression. The red dashed line represents the fitted linear regression obtained using ordinary least squares and each point corresponds to a bladder cancer biopsy (n=73, R²=0.35, p<0.0001).

### TFEB and HGS/Hrs regulate endosomal membrane fluidity

Integrins are transmembrane adhesion proteins whose trafficking dynamics are strongly influenced by membrane properties such as lipid order and membrane fluidity (Moreno-Layseca et al. 2019; Lietha and Izard 2020). Membrane fluidity and lipid order can be imaged using fluorescent probes sensitive to polarity (solvatochromic dyes), such as derivatives of Laurdan (Parasassi et al. 1991; Owen et al. 2011) and Nile Red (Kucherak et al. 2010; Klymchenko 2023). We thus tested whether TFEB and HGS/Hrs could control cellular membrane properties by evaluating emission of the Laurdan fluorescent dye with a emission shift from blue to red in less packed lipids with increased water penetration (Orlikowska-Rzeznik et al. 2023). Ratiometric imaging of whole cells incubated with Laurdan showed that TFEB or HGS/Hrs depletion increased red/blue ratio in both KU19-19 and JMSU1 cells (**Fig. 4A,C; SFig. 2A,C**, respectively) indicating an increase in water penetration and membrane fluidity. Contrary, cells that exogenously overexpressed TFEB-EGFP or HGS-EGFP sowed reduced red/blue ratio indicative of more ordered/stiff membranes (**Fig. 4B,D**, for KU19-19 cells and **SFig. 4B,D** for JMSU1 cells), with a particularly strong effect seen for overexpressed HGS/Hrs. Importantly, HGS/Hrs overexpression markedly reduced the red/blue ratio in TFEB-depleted cells, restoring membrane order despite TFEB loss (**Fig. 4D** for KU19-19 cells and **SFig. 4D** for JMSU1 cells). These findings place HGS/Hrs downstream of TFEB in the pathway controlling endosomal membrane fluidity.

**Figure 4.**
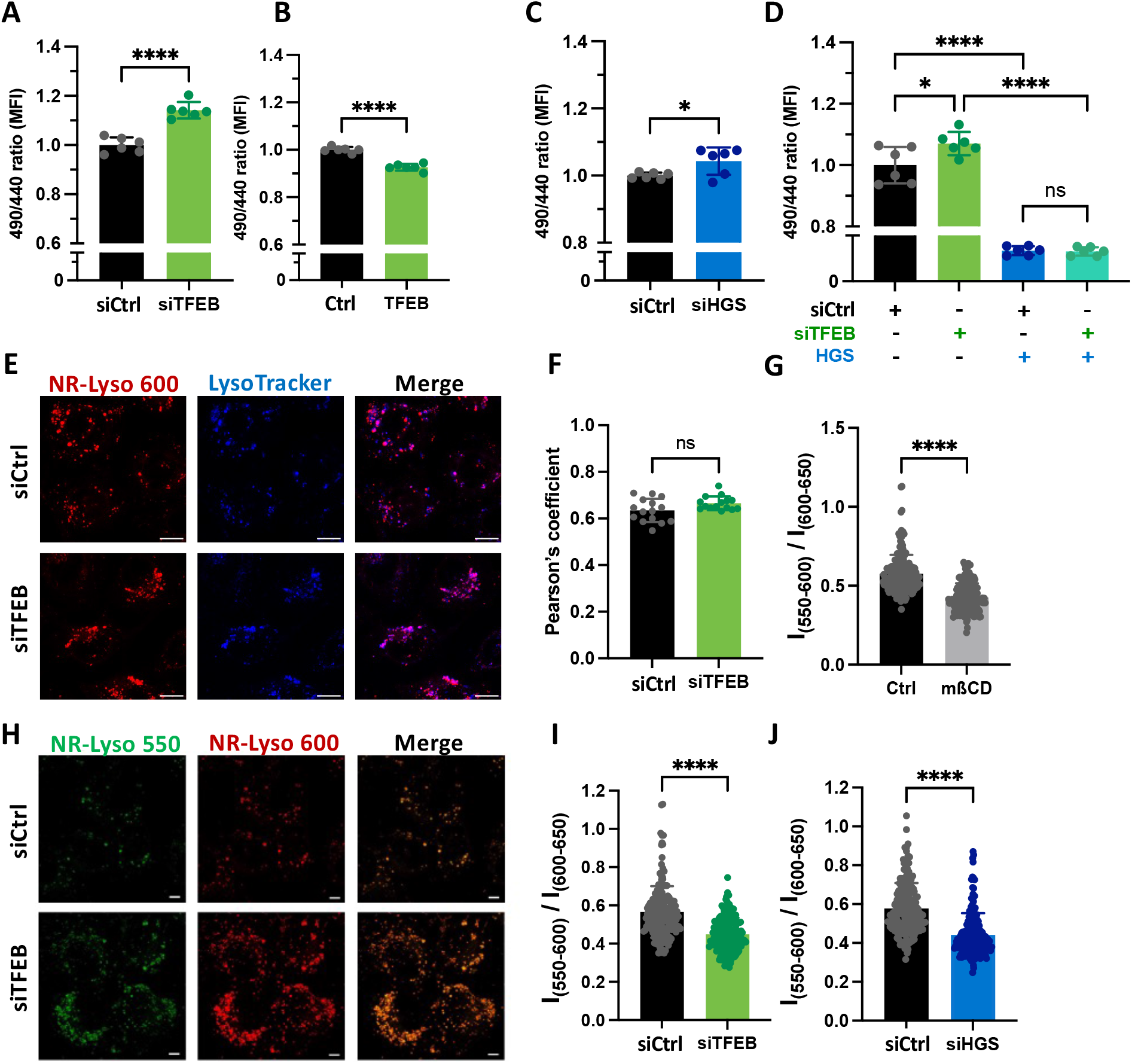
TFEB and HGS/Hrs regulate endosomal membrane fluidity. (A) Ratiometric measurement of membrane fluidity by flow cytometry using the polarity-sensitive Laurdan dye in KU19-19 cells 72 h post-transfection with siControl (Ctrl) or siTFEB. Data represent mean ± SD of N=3; ****p < 0.0001 (unpaired Student’s t test with Welch’s correction). (B) Ratiometric measurement of membrane fluidity by flow cytometry using the polarity-sensitive Laurdan dye in KU19-19 cells 24 h post-transfection with the EGFP vector (Ctrl) or pEGFP-TFEB plasmid (TFEB). Data represent mean ± SD of N=3; ****p < 0.0001 (unpaired Student’s t test with Welch’s correction). (C) Ratiometric measurement of membrane fluidity by flow cytometry using Laurdan dye in KU19-19 cells 48 h post-transfection with siHGS or siCtrl. Data represent mean ± SD of N=3; *p < 0.05 (unpaired Student’s t test with Welch’s correction). (D) Ratiometric measurement of membrane fluidity by flow cytometry using Laurdan dye in KU19-19 cells 72 h post-transfection with siTFEB or siCtrl and 24 h post-transfection with the pCS2-HRS-EGFP plasmid (HGS) or EGFP vector (Ctrl). Data represent mean ± SD of N=3; n.s. not significant; *p < 0.05; ****p < 0.0001 (unpaired Student’s t test with Welch’s correction). (E) Fluorescence staining of NR-Lyso (red) and LysoTracker (blue) in KU19-19 cells treated with siCtrl or siTFEB. Scale bars, 10 μm. **(F)** Quantification of Pearson’s correlation coefficient for NR-Lyso and LysoTracker colocalization in KU19-19 cells transfected with siCtrl or siTFEB (arbitrary units, a.u.). Data represent mean ± SD from N=3; ns not significant (Student’s t test). (G) Quantification of the green/red fluorescence intensity ratio (I₅₅₀–₆₀₀/I₆₀₀–₆₅₀) of the NR-Lyso probe in KU19-19 cells treated for 30 min with 10 mM methyl-β-cyclodextrin (mβCD). Data represent mean ± SD of N=3; ****p < 0.0001 (Mann– Whitney U test). (H) Representative ratiometric confocal images of KU19-19 cells stained with NR-Lyso. Scale bars, 10 μm. (I) Quantification of the green/red fluorescence intensity ratio (I₅₅₀–₆₀₀/I₆₀₀–₆₅₀) of the NR-Lyso probe in KU19-19 cells transfected with siCtrl or siTFEB. Data represent mean ± SD of three independent experiments; ****p < 0.0001 (Mann–Whitney U test). (J) Quantification of the green/red fluorescence intensity ratio (I₅₅₀–₆₀₀/I₆₀₀–₆₅₀) of the NR-Lyso probe in KU19-19 cells transfected with siCtrl or siHGS. Data represent mean ± SD of N=3; ****p < 0.0001 (Mann–Whitney U test).

Then, we investigated membrane properties of acidic endosomes that are under control of TFEB using the Red Nile Lyso (NR-Lyso), an endosomal membrane polarity probe (Danylchuk et al. 2021). Similarly to Laurdan, Nile Red fluorophore is sensitivity to lipid order and membrane fluidity by shifting its emission band (Klymchenko 2017), while the targeting group of NR-Lyso ensures specific localization in the dye inside endosomal membranes. NR-Lyso accumulated in acidic compartments in both cell lines, and its co-localization with the lysotracker probe was unchanged upon TFEB KD (**Fig. 4E,F; SFig. 4E,F**). We validated NR-Lyso sensitivity to membrane fluidity/lipid order by treating cells with methyl-β-cyclodextrin (mβCD) that extracts cholesterol from membranes. As expected, ratiometric analysis of NRLyso revealed a significant fluorescence shift toward longer wavelengths following mβCD treatment (**Fig. 4G, SFig. 4G**), consistent with increased endosomal membrane fluidity. A comparable shift was observed in both cell lines following TFEB depletion (**Fig. 4H,I, SFig. 4H,I**) or HGS/Hrs depletion (**Fig. 4J, SFig. 4J**), indicating that both proteins contribute to the maintenance of endosomal membrane order. Together, these findings establish TFEB and HGS/Hrs as regulators of the biophysical state of endosomal membranes.

### TFEB regulates cellular lipid composition

Because membrane fluidity is intrinsically linked to lipid composition (Sezgin et al. 2017), we next performed lipidomic analysis of control JMSU1 and TFEB depleted cells. We identified significant alterations in lipid composition following siTFEB treatment (**Fig. 5A**), particularly a significant decrease (p < 0.01) in cholesterol, which accounts for about 20 pmol% of lipids in these cells. Filipin staining in control and siTFEB cells confirmed a decrease in cellular cholesterol levels upon TFEB KD in both bladder cancer cell lines (**Fig. 5B, SFig. 5A**). Moreover, ether-phosphatidylcholine (PCO) showed significant reduction (p < 0.01). PCO is known to increase membrane order by increasing lipid packing due to the sn-1 ether bond. It goes in line with a more disordered membrane state in TFEB depletion (**Fig. 5A**). PCO accounts for about 2,5 pmol% of lipids and are known as plasmalogen species that are implicated in membrane structure. At the same time, triacylglycerols (TAGs) that account for about 2 pmol% of lipids, significantly increased (p < 0.01) (**Fig. 5A**). Visualization of endogenous lipid droplets with bodipy did however not show any changes in either total lipid droplet fluorescence or lipid droplet numbers (**Fig. 5C-E**, **SFig. 5B,C**). Analyses of the overall acyl chain profile of TAGs, which represents the fraction of each lipid species within a given lipid class, revealed that TFEB depletion particularly increased the presence of triacylgrycerides of longer than 46 carbon chains (**Fig. 5F**). Moreover, sphingolipids of the glucosylceramide (GlcCer) and globotriaosylceramide (Gb3) classes, accounting to about 4 and 1,5 pmol % of lipids, respectively, were significantly increased (p < 0.01) (**Fig. 5A**). We further analyzed the degree of unsaturation of abundant phospholipids (> 1 mol%). We found that phospholipids (PC, PE, PEO, PI, PS) revealed a significant decrease of mono- and di-unsaturated short fatty acid (<C36) accommodated with a significant increase in polyunsaturated long fatty acid (>C36) (**SFig. 5D, E**) upon siTFEB. For TAGs, we found a particular increase of polyunsaturated fatty acids of more than 6 double bounds and longer than C58/C52 (**Fig. 5G, SFig. 5F**). For sphingolipids such as Gb3, Gb4 and GlcCer, we detected a total increase in mono- and di-unsaturated short fatty acid (**SFig. 5F,G**). No significant changes in fatty acid saturation were found in DAG, cholesterol esters, ceramide, GM2 and LacCer. Thus, TFEB depletion resulted in a significant alteration in membrane lipid composition on the whole cell level, particularly decrease of cholesterol accompanied by the accumulation of poly-unsaturated long chain fatty acid, and mono-unsaturated sphingolipids that support the observed membrane fluidity alterations in siTFEB conditions.

**Figure 5.**
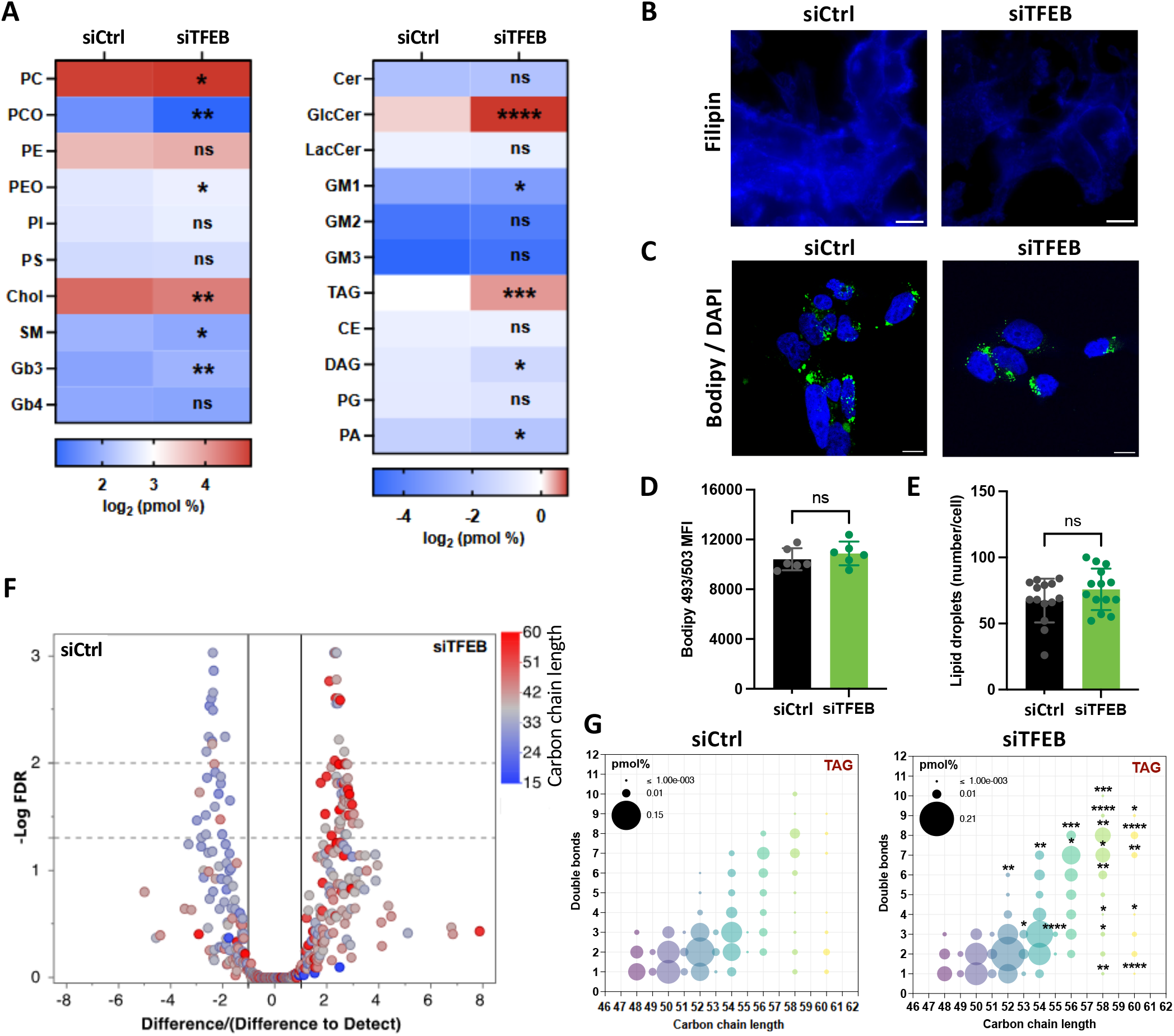
TFEB regulates lipid composition in JMSU1 cells. (A) Total lipids were extracted from JMSU1 cells 72 h post-transfection with siControl (Ctrl) or siTFEB, followed by quantification of phospholipid species and total cholesterol. Heatmap shows relative/absolute changes of lipid classes. Data are expressed as log₂-transformed lipid quantities (pmol %). Significance was determined by an unpaired Student’s t test with Welch’s correction; ns, not significant; *p < 0.05; **p < 0.01; ***p < 0.001; ****p < 0.0001. (B) Representative confocal images of Filipin fluorescence in control and TFEB-depleted JMSU1 cells 72 h post-transfection, stained with Filipin. Scale bars, 10 μm. (C) Representative confocal images of control and TFEB-depleted JMSU1 cells 72 h post-transfection, stained with Bodipy 493/503 (green) and counterstained with DAPI. Scale bars, 10 μm. (D) Flow cytometry analysis of lipid droplet content in control and TFEB-depleted JMSU1 cells stained with Bodipy 493/503 (green) 72 h post-transfection. Data represent mean ± SD of N=3; ns, not significant (unpaired Student’s t test with Welch’s correction). (E) Quantification of lipid droplet number per cell in JMSU1. Data represent mean ± SD of N=3; ns, not significant (Mann–Whitney U test). (F) Volcano plot showing differential abundance of total carbon chain lengths of lipid species between siTFEB-transfected and control cells. The ratio “Difference / Difference to detect” indicates the difference in total chain length between conditions normalized by the minimum detectable difference. . Longer and shorter fatty acids are shown in red and blue, respectively. (G) Bubble plot of TAG species in control (upper panel) and TFEB-depleted (lower panel) JMSU1 cells. Each circle represents a TAG species defined by its number of carbon atoms and double bonds. Circle size indicates the average quantity of each species, and fill color represents total carbon content. Significance was determined by two-way ANOVA with Šídák’s multiple comparisons test; *p < 0.05; **p < 0.01; ***p < 0.001; ****p < 0.0001.

### Endosomal membrane fluidity regulates endosomal positioning and ITGβ5 trafficking

Finally, we tested if altered membrane properties could regulate endosomal organization and thereby influence integrin trafficking. Cellular membrane fluidity can be instructed by addition of exogenous lipid species (Bergen et al. 2023). We therefore treated KU19-19 cells for 24 h with either the monounsaturated oleic acid (C18:1) or the saturated palmitic acid (C16:0) and assessed endosomal membrane fluidity using the NR-Lyso probe. Consistent with previous studies (Ehringer et al. 1990; De Santis et al. 2018; Manni et al. 2018; Harayama and Antonny 2023), oleic acid significantly increased endosomal membrane fluidity, whereas palmitic acid had no detectable effect (**Fig. 6A,B**).

**Figure 6.**
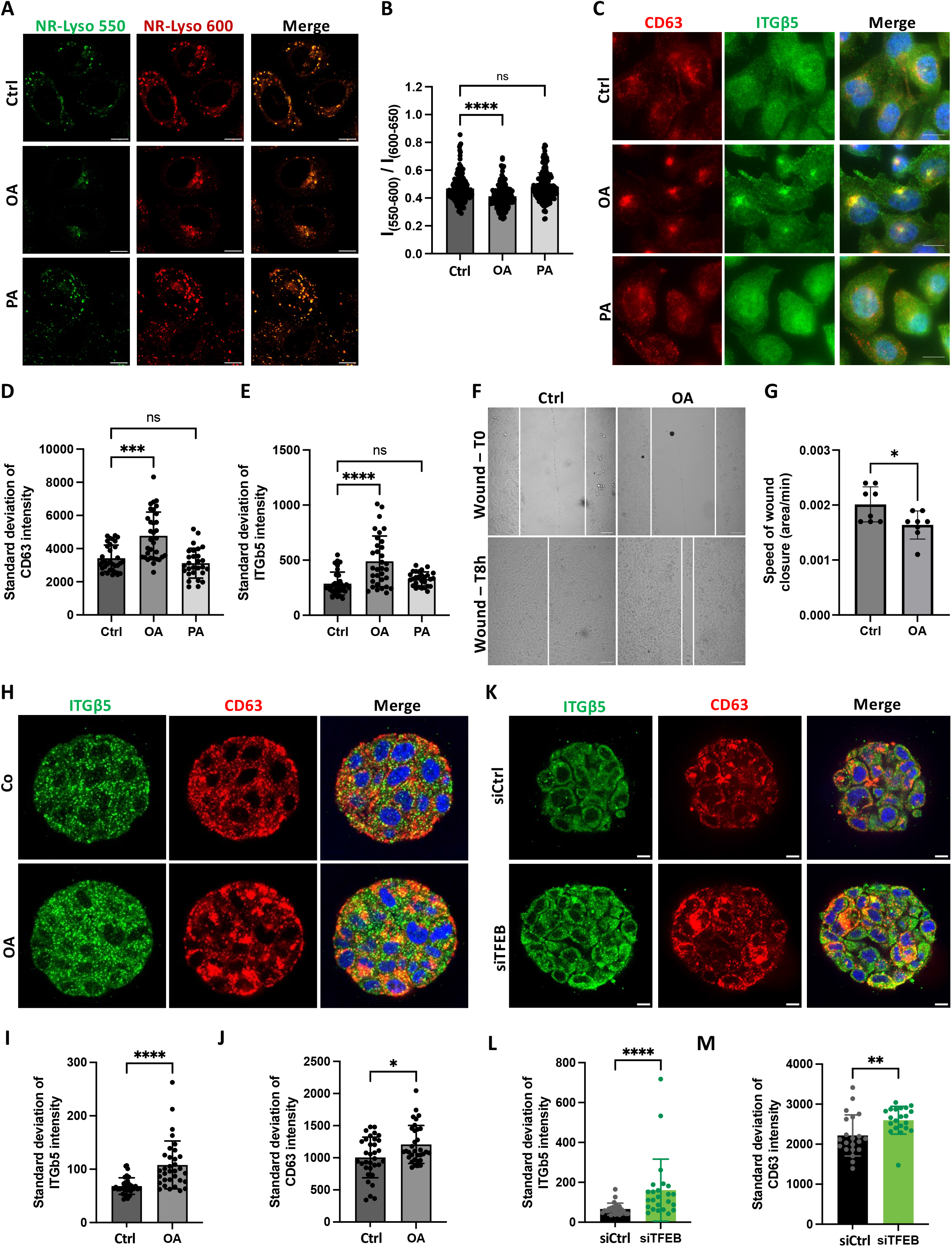
Fatty acid–regulated endosomal membrane fluidity controls MVB positioning and ITGβ5 trafficking. (A) Representative ratiometric confocal images of NR-Lyso-stained KU19-19 cells treated for 24 h with ethanol 100% (Ctrl), 50 µM oleic acid (OA), or palmitic acid (PA). (B) Quantification of the green/red fluorescence intensity ratio (I₅₅₀–₆₀₀ / I₆₀₀–₆₅₀) in KU19-19 cells treated for 24 h with ethanol 100% (Ctrl), 50 µM oleic acid (OA), or palmitic acid (PA). Data represent mean ± SD of N=3; ns, not significant, ****p < 0.0001 (Kruskal–Wallis test followed by Dunn’s multiple comparisons). (C) Confocal images of KU19-19 cells treated for 24 h with ethanol 100% (Ctrl), 50 µM oleic acid (OA), or palmitic acid (PA) and immunostained for CD63 (red) and ITGβ5 (green). Nuclei were counterstained with DAPI. Scale bars, 10 μm.Scale bars, 10 μm. **(D)** Quantification of CD63 standard deviation intensity of KU19-19 cells treated for 24 h with ethanol 100% (Ctrl), 50 µM oleic acid (OA), or palmitic acid (PA). Data represent mean ± SD of N=3; n.s. not significant; ***p < 0.001 (unpaired Student’s t test with Welch’s correction). **(E)** Quantification of ITGβ5 standard deviation intensity of KU19-19 cells treated for 24 h with ethanol 100% (Ctrl), 50 µM oleic acid (OA), or palmitic acid (PA). Data represent mean ± SD of N=3; n.s. not significant; ****p < 0.0001 (unpaired Student’s t test with Welch’s correction). **(F)** Representative wound-healing assay of KU19-19 cells in the presence of either ethanol 100% (Ctrl) or oleic acid (OA). Wound closure was monitored over 360 min (8 h). Scale bars, 100 μm **(G)** Quantification of wound closure in the presence of ethanol 100% (Ctrl) or oleic acid (OA), calculated as |Tₙ–T₀|/T₀ in area/mins. Data represent mean ± SD of N=2; ****p < 0.0001 (Student’s t test). (H) KU19-19 cells grown for 24 h in soft hydrogel–containing microcavity arrays were treated with 100 µM OA for 24 h, followed by immunostaining for CD63 (red) and ITGβ5 (green) and imaging by spinning disk microscopy. Nuclei were counterstained with DAPI. Scale bars, 10 μm. (**I,J**) Quantification of ITGβ5 (I) and CD63 (J) standard deviation intensity from images obtained in (H). Data represent mean ± SD of N=3; *p < 0.05; ****p < 0.0001 (unpaired Student’s t test with Welch’s correction). (M) KU19-19 cells transfected with siTFEB for 24 h were grown in microcavity arrays for 48 h, followed by immunostaining for CD63 (red) and ITGβ5 (green) and imaging by spinning disk microscopy. Nuclei were counterstained with DAPI. Scale bars, 10 μm. (**L,M**) Quantification of ITGβ5 (L) and CD63 (M) standard deviation intensity from images obtained in (K). Data represent mean ± SD of N=3; **p < 0.01; ****p < 0.0001 (unpaired Student’s t test with Welch’s correction).

Strikingly, oleic acid treatment induced an obvious clustering of CD63-positive endosomes, whereas palmitic acid did not (**Fig. 6C)**, indicating that membrane fluidity influences endosomal positioning. We next examined whether ITGβ5 trafficking was sensitive to changes in membrane fluidity upon addition of exogenous lipid species. Immunofluorescence analysis revealed a marked accumulation of ITGβ5 within clustered CD63-positive endosomes following oleic acid treatment, whereas palmitic acid had no effect (**Fig. 6C**). Quantification of the clustering of CD63 and ITGβ5 using the standard deviation of fluorescence intensity, showed that oleic acid treatment resulted in significantly higher values indicating a more heterogeneous and spatially concentrated signal (**Fig. 6D,E**). Consistent with these trafficking defects, oleic acid significantly reduced wound closure speed in KU19-19 cells (**Fig. 6F,G**). Together, these findings demonstrate that increasing endosomal membrane fluidity is sufficient to induce ITGβ5 accumulation within MVBs and impair cell migration. To further test the relationship between membrane fluidity and TFEB-dependent phenotypes, we supplemented siCtrl and siTFEB cells with cholesterol, which decreases membrane fluidity. As expected, cholesterol treatment significantly reduced the fluidity of acidic endosomes irrespective of TFEB status (**SFig. 6A-D**). Consistent with this effect, cholesterol reduced the clustering of both CD63-positive endosomes and ITGβ5 in TFEB-depleted cells (**SFig. 6C**), indicating that restoration of membrane order rescues the endosomal organization defects associated with TFEB loss.

To assess the relevance of these findings in a three-dimensional context, we examined ITGβ5 distribution in KU19-19 tumoroids grown in bioengineered compliant microcavities (Ouni et al. 2025). Clustering was quantified using the standard deviation of fluorescence intensity, with higher values indicating a more heterogeneous and spatially concentrated signal. Oleic acid treatment significantly increased clustering of ITGβ5 within CD63-positive MVBs (**Fig. 6G-I**). Similarly, TFEB depletion promoted clustering of both CD63-positive structures and ITGβ5 in 3D cultures (**Fig. 6K-M**). Taken together these results indicated that endosomal membrane fluidity can regulate endosomal positioning and ITGβ5 trafficking during 2D cell migration and in 3D culture, linking membrane biophysical state to integrin trafficking and cell migration.

## Discussion

Our study uncovers a previously unrecognized connection between TFEB and integrin trafficking by identifying endosomal membrane fluidity as the mechanistic bridge between TFEB transcriptional control and cell migration. Although elevated TFEB expression has been associated with migratory phenotypes in non–small cell lung cancer and prostate cancer (Giatromanolaki et al. 2015; Zhu et al. 2021), the molecular basis for this correlation has remained unresolved. Here, we demonstrate that TFEB governs the biophysical state of endosomal membranes and that these physical properties, in turn, determine the intracellular routing of integrins, particularly ITGβ5 in bladder cancer cells (**SFig. 6E**). This framework provides a mechanistic rationale for TFEB-dependent regulation of migration.

Integrin sorting is orchestrated within the endosomal network, where activation state and adaptor engagement determine trafficking fate (Chastney et al. 2025). Although ITGβ5 trafficking remains poorly characterized, the intracellular trafficking routes of ITGβ1 have been extensively studied. After endocytosis, active ITGβ1 is recycled directly to the plasma membrane through actin- and WASH-dependent mechanisms (MacDonald et al. 2018), whereas inactive ITGβ1 traffics via the trans-Golgi network in a retromer-dependent route before re-secretion (Shafaq-Zadah et al. 2016). Alternatively, ITGβ1 may be targeted to multivesicular bodies (MVBs) for lysosomal degradation through ubiquitination and ESCRT-dependent sorting (Lobert et al. 2010; Yu et al. 2024). Although we did not detect changes in total ITGβ5 levels upon TFEB depletion within the experimental time frame, our data indicate pronounced endosomal retention of ITGβ5, reminiscent of phenotypes observed upon ESCRT-0 disruption for ITGβ1, where integrin trapping correlates with impaired directional migration and defective focal adhesion turnover (Lobert et al. 2010; Lobert and Stenmark 2012; Matthew-Onabanjo et al. 2020). Future work will need to determine in more detail how trafficking routes of ITGβ5 are regulated by the membrane states of different endosomes.

We demonstrate that increasing endosomal membrane fluidity, either by oleic acid supplementation (Ehringer et al. 1990; De Santis et al. 2018; Manni et al. 2018) or by depletion of the ESCRT-0 subunit HGS/Hrs, is sufficient to induce perinuclear clustering of multivesicular bodies (MVBs) and impair ITGβ5 trafficking and cell migration. Conversely, decreasing membrane fluidity through cholesterol supplementation rescues the phenotypes associated with TFEB depletion. Membrane fluidity, defined as the inverse of viscosity and governed by lipid packing and molecular interactions, controls protein partitioning. Consistent with prior work showing that membrane physical properties regulate integrin trafficking (Lietha and Izard 2020; Mikhajlov et al. 2025), either via transmembrane domain sorting or adaptor-mediated mechanisms (Calderwood et al. 2013; Ge et al. 2018; Lietha and Izard 2020), our findings place lipid-driven biophysical regulation upstream of adhesion receptor routing.

Mechanistically, we identify HGS/Hrs as a key intermediate linking TFEB to endosomal membrane organization. TFEB depletion reduces HGS/Hrs levels, and HGS/Hrs loss is sufficient to increase endosomal membrane fluidity, establishing it as a downstream regulator of endosomal biophysical state. Conversely, HGS/Hrs overexpression restores endosomal distribution and reverses the perinuclear accumulation of ITGβ5. HGS/Hrs is a FYVE-domain–containing protein that binds PI3P-enriched endosomes (Gaullier et al. 1998; Raiborg, Bremnes, et al. 2001), and nucleates ESCRT-0 assemblies to concentrate ubiquitinated cargo for their sorting into ILV at MVBs (Raiborg, Grønvold Bache, et al. 2001). Notably, our previous work has shown that TFEB regulates PI3P levels on endo-lysosomes in bladder cancer cells by transcriptionally regulating expression of VPS34 (PIK3C3) (Mathur et al. 2023), the lipid kinase responsible for PI3P production (Gambardella et al. 2020; Mathur et al. 2023); thus providing a plausible axis through which TFEB controls HGS/Hrs function. Strikingly, we did not observe alterations in downstream ESCRT components such as TSG101 or ALIX, suggesting that the observed phenotype may reflect ESCRT-0– specific functions independent of canonical ESCRT progression.

Emerging evidence indicates that ESCRT-0 components can form biomolecular condensates that organize membrane domains. Analogous to the plant FYVE-domain protein FREE1, which mediates membrane bending and intraluminal vesicle formation via condensate-driven modulation of membrane line tension (Wang Y. et al. 2024), HGS/Hrs has been shown to form 2D condensates on negatively charged membranes that assemble flat clathrin lattices (Hakala et al. 2026). Cholesterol promotes HGS condensate formation and accumulates beneath HGS–clathrin microdomains (Hakala et al. 2026). We therefore speculate that HGS/Hrs condensates locally trap cholesterol to generate membrane subdomains of reduced fluidity. Consistent with this model, HGS depletion increased endosomal membrane fluidity, potentially reflecting diminished cholesterol confinement. In addition to altering membrane properties, HGS depletion reduced levels of the small GTPase RAB27, a regulator of MVB anterograde trafficking (Ostrowski et al. 2010). These findings suggest that ESCRT-0 components coordinate membrane composition with organelle positioning and motility, linking membrane biophysics to intracellular transport dynamics.

Beyond its established role in lysosomal biogenesis, our work identifies TFEB as a regulator of lipid composition and membrane fluidity in bladder cancer cells. TFEB depletion reduced cholesterol levels and increased the presence of poly-unsaturated fatty acid chains in phospholipids that is consistent with increased membrane fluidity (Veatch and Keller 2002). We found that membrane fluidity was increased at acidic endosomes as well as the whole cell level in the bladder cancer model. In plasma membranes, lipid order has been proposed as a crucial characteristic of membrane microdomains, which define membrane organization and regulate the function of membrane proteins (Lingwood and Simons 2010). The role of membrane fluidity in acidic endosomes is less explored, even though it is well established that endosomal maturation is associated with a decrease in cholesterol content and thus an increase in membrane fluidity (Kobayashi et al. 1999; Huotari and Helenius 2011; Darwich et al. 2014). TFEB targets include enzymes involved in sphingolipid metabolism, and we observed alterations in glycosphingolipid species, further implicating TFEB in controlling membrane composition. How these lipid changes intersect with peroxisomal metabolism (Evans et al. 2019; Mao et al. 2022; Sass et al. 2021; Settembre et al. 2013), lipid droplet homeostasis (Menon et al. 2023), or membrane saturation sensors (Covino, Hummer, and Ernst 2018; Jian et al. 2022; Ruiz et al. 2019; Y.-T. Wang et al. 2019) remain important questions for future studies.

Interestingly, TFEB activation in aggressive bladder cancer cells does not robustly induce canonical lysosomal biogenesis programs (Mathur et al. 2023), consistent with emerging evidence that TFEB functions are context dependent (Calcagnì et al. 2016; Doronzo et al. 2019; Kim et al. 2021). Depending on cell type, TFEB appears to preferentially influence distinct endolysosomal compartments, early endosomes, MVBs, or lysosomes (Mathur et al. 2023; Palmieri et al. 2011; Settembre et al. 2011), suggesting that its core function may extend beyond simple biogenesis to active maintenance of endomembrane homeostasis. We speculate that membrane remodeling downstream of TFEB could induce lysosome biogenesis and autophagy as a terminal phenotype for lipid detoxification (Jain and Zoncu 2026; Radulovic et al 2026; Thelen and Zoncu 2017). In bladder cancer cells, this stage seems to not be reached yet, giving rise to a milder phenotype of cell migration.

We propose that the primary function of TFEB in this context is the maintenance of endosomal membrane homeostasis. By transcriptionally coordinating lipid metabolism and ESCRT-0 components, TFEB establishes cholesterol-enriched membrane microdomains at acidic endosomes that constrain membrane fluidity within MVBs. These biophysical properties function as a regulatory layer instructing integrin sorting decisions that affect cellular migration and adhesion. Our results substantialize previous observations that endosomal sorting relies on cell lipid homeostasis (Alonso-Bivou et al. 2025; Carpentier et al. 2025; Otakhor et al. 2025; Peng et al. 2025; Song et al. 2022). In this framework, membrane material state is not merely a consequence of lipid metabolism but an instructive determinant of trafficking outcomes. Because the endosomal network defines the trafficking, and thus, plasma membrane exposure of cell surface receptors and adhesion molecules, transcriptional control of membrane material state can directly reshape cell adhesion and migratory behavior.

Together, our study reframes TFEB as a regulator of membrane properties in bladder cancer cells, and positions endosomal fluidity as a critical determinant of integrin trafficking and cell migration. By linking transcriptional control of lipid metabolic programs to membrane properties and organization of the endosomal system, we provide a conceptual framework in which metabolism, membrane material state, and cell behavior are mechanistically integrated.

## Material and Methods

### Cell culture and reagents

Bladder cancer cell lines KU19-19 and JMSU1 were obtained from the American Type Culture Collection (ATCC). KU19-19 cells are derived from a T3b-stage muscle invasive bladder carcinoma (MIBC) and JMSU1 are derived from a T4-stage MIBC. Cells were grown in a 5% CO_2_ humidified atmosphere at 37°C in RPMI 1640 (Thermo Fisher Scientific) supplemented with 10% SVF and 100 units/mL of penicillin/streptomycin. Cells were split twice a week and media was changed 24 h before harvesting the cell with fresh complete RPMI 1640 media. Reagents used are presented in **Supplementary Table S1**.

### siRNA transfection

KU19-19 and JMSU1 cells were seeded in 6-well plates at a density of 100 000 cells/well and 200 000 cells/well, respectively. After 24 h of seeding, cells were transfected with either a mixture of four or individual predesigned siRNAs targeting *TFEB* (ON-TARGETplus human, L-009798-00-0005, Dharmacon) or *HGS* (ON-TARGETplus human, L-016835-00-0005, Dharmacon) or a negative control siRNA targeting Luciferase (5’-CGTACGCGGAATACTTCGA-3’, Sigma Aldrich) at a final concentration of 50 nM. Briefly, in a 1,5-mL tube, 980 μL of Opti-MEM medium (Thermo Fisher Scientific) were mixed with 20 μL of lipofectamine RNAimax (Invitrogen). In a second tube, 496 μL of Opti-MEM medium were mixed with 4 µL of the siRNA pool at 50µM. After 5 minutes of incubation, the two tubes were mixed. The reaction mixture was incubated for 20 min at room temperature and dispensed drop by drop onto the cells. Cells were incubated with siRNAs targeting *TFEB* for 72 h or *HGS* for 48 h before subsequent analysis.

### RNAseq analysis

RNA sequencing was performed by the Minos Bioscience startup (ESPCI, PSL University) on KU19-19 cells transfected either with 72-h siTFEB or 72-h siLUC. In brief, genes with ≥10 total counts and at least 1 count in each condition were retained to reduce dropout effects. Counts were normalized to counts per million (CPM) to account for library size. Differential expression was assessed using log₂ fold changes between conditions. Gene lists were analyzed for Gene Ontology (GO) enrichment using the R package clusterProfiler and topGO, following standard over-representation analysis workflows. Similar analysis were also performed for gene lists obtained from (Diamant et al. 2025; Gambardella et al. 2020; Palmieri et al. 2011).

### RT-qPCR

Total cellular RNA was extracted using TRIzol reagent (Thermo Fisher Scientific). RNA concentrations were determined using NanoDrop 2000 spectrophotometer (Thermo Fisher Scientific). 1 µg RNA was reverse-transcribed using M-MVL reverse transcriptase, according to the manufacturer’s protocol (Thermo Fisher Scientific). Thereafter, cDNA was used in real-time PCR with the KAPA SYBR FAST (Sigma Aldrich). The relative expression levels of the mRNA of interest were determined by the 2^-ΔCt^ method and normalized to the expression levels of human actin (*ACTB)*. The primer sequences used are listed in **Supplementary Table S2**.

### Wound healing assay

After a 48-h siHGS or 72-h siTFEB transfection, KU19-19 cells were transferred to a glass bottom plate precoated with fibronectin (concentration at 50μg/mL). When indicated, 10μM cilengitide (CLG), 300 ng/μL mitomycin c (MMC), or DMSO as control was added. Before imaging, a scratch was made using a tip on the cell layer and media was changed to remove scratched cells. Cells were imaged using a 10X objective on Leica Thunder Microscope (Leica Microsystems) or Videomicroscope (Leica Videomicroscope with SP8 Stand). Images were taken every 10 min for 6 h and during the acquisition the temperature is maintained at 37°C and the CO_2_ at 5%. Quantification was performed by measuring the cell-free area each 10 frames using the Fifi software. The areas were normalized by the T0 frame for each conditions following the formula abs((Tn-T0)/T0). The normalized areas were plotted over time, and the slope was used for the quantification.

### Single cell migration assay

After a 48-h siHGS or 72-h siTFEB transfection, JMSU1 cells were transferred to glass bottom plate precoated with fibronectin (concentration at 50μg/mL). Cells were imaged using a 10X objective on the Leica Thunder Microscope or Leica SP8 Stand). Images were taken every 10 min for 6 h. Cell tracking was performed using the Manual Tracking plugin of the Fiji software. In short, single cell were tracked on every frame for 6 hours and the displacement of the cell between two frames (10 minutes) was calculated using the plugin. The mean speed for each cell was plotted.

### Adhesion assay

Cells were detached from the plate using Versen (Thermofisher) and allowed for 1 h to adhere on fibronectin-coated coverslips (concentration at 50 μg/mL). Then, cells were washed three times in PBS 1X and fixed in 4% PFA. Cells were permeabilized with 0,005% saponin and 0,2% BSA, followed by staining with deep red cell mask (A57245, Thermo Fisher Scientific) and DAPI (62248, Thermo Fisher Scientific). Coverslips were washed twice in PBS 1X and mounted on slides using Fluoromount-G (Thermo Fisher Scientific). Images were acquired with the Leica Thunder Microscope.

### Immunofluorescence staining

Cells were seeded 24 h before fixation in 24-well plates containing sterilized fibronectin precoated glass coverslips. Cells were fixed in 4% PFA for 20min at room temperature, washed three times in PBS 1X and blocked/permeabilized with PBS 1X supplemented with saponin 0,05% and BSA 0,2% (PBS+) for 20 min at room temperature. Cells were then incubated with primary antibodies diluted in PBS+ buffer (**Supplementary Table S3**) for 1 h at room temperature, followed by washing with PBS 1X and incubation with DAPI (62248, Thermo Fisher Scientific) and secondary antibodies diluted in PBS+ buffer, donkey anti-mouse Alexa Fluor 488 (715-545-151, Jackson Immuno Research) and anti-rabbit Alexa Fluor Cy3/568 (711-165-152, Jackson Immuno Research), used at 1:400 for 45 min at room temperature in the dark.

For ITGb5 staining or double staining of CD63/ITGb5, cells were seeded in sterilized glass coverslips previously coated with fibronectin for 1 h (1:20 in water, F1141, Sigma). After treatment, cells were fixed in 4% PFA for 20min at room temperature, washed three times in PBS 1X and permeabilized with PBS 1X supplemented with 0,1% Triton X-100 for 20 min at room temperature. After 1 h of saturation at room temperature with PBS 1X supplemented with 0,2% BSA, cells were then incubated with primary antibodies (**Supplementary Table S3**) in saturation buffer overnight at 4°C. After staining, cells were washed and incubated with secondary antibodies diluted in saturation buffer for 45 min at room temperature. Isotype-matched IgG antibodies were used as negative control. Coverslips were then washed three times and mounted with Fluoromount-G (Thermo Fisher Scientific) on glass slides. After drying overnight at 4°C, image acquisition was performed using Leica SP8 confocal microscope or Spinning Disk NR1.

### Plasmid transfection

On the 2nd day following siRNA transfection and 48h before imaging in a 12-well plate, cells were transfected with 2µg of clathrin-adaptor β2-GFP plasmid with 4µL of PEI in 100µL of Opti-MEM media, according to the manufacturer’s instructions (Polyplus). Briefly, reagents were vortexed and incubated 20 min at room temperature before being added to the cells drop by drop. Medium was changed for complete RPMI medium 4 h post-transfection.

### TIRF microscopy

Cells were plated on a fibronectin coated glass bottom 35mm petri dish (ibidi). Dynamics of β2-GFP was acquired by imaging cells every 5 seconds for 5 minutes using the TIRF mode of Gataca based onTi2 Eclipse Nikon. Data analysis was performed using TrackMate plugin of the Fiji software.

### STED

Cells were washed with cold PBS 1X, fixed in 100% cold methanol for 20 min on ice and permeabilized with PBS1X supplemented with 0,01% Triton X-100 20 min at room temperature. Cells were blocked using 0,2% BSA for 1 hour and then incubated for 1 h at room temperature with primary antibodies (**Supplementary Table S3**). After 3 rinses for 5 min in PBS 1X, STED secondary antibody was incubated at 1:100 in PBS 1X + 0,2% BSA for 1 h at room temperature. After 3 rinses for 5 min in PBS 1X, cells were rinsed in water, then mounted on a slide using 5µL of STED montage medium (Thermo Fisher Scientific). After drying overnight at 4°C, images were taken using the Abberior Stedycon on Leica SPE base.

### Exosome isolation and quantification

Exosomes isolation and detection experiments were performed on the CurieCoreTech-Extracellular Vesicles Platform (Curie Institute, Paris, France). In brief, cells were plated in two petri plates (B10) and transfected with siRNA against TFEB as previously described. On the 2^nd^ day, cells were transferred in T175 flask for amplification, washed with PBS 1X to remove lipidic or protein particles from serum, and fresh medium without serum was added. Cells were incubated for 24 hours for extracellular vesicles (EVs) productions. On the EVs isolation day, EV-containing supernatant was collected and centrifuged at 350 g for 10 min at 4°C to eliminate dead cells. EV-producing cells were collected to assess cellular viability using trypan blue and proteins were extracted as control. The supernatant was centrifuged twice at 350g during 10 min at 4°C and once at 2 000 g for 20 min at 4°C to remove large particles including cell debris. Supernatant was transferred into ultracentrifuge tubes (SW32) and centrifuged at 10 000 g for 30 min at 4°C to remove large extracellular particles. Supernatant was transferred to new ultracentrifuge tubes and ultracentrifuged at 100 000 g for 90 min at 4°C. The pellet was washed and resuspended in 50 μL in 0,22 μm filtered PBS 1X. Samples were divided into two parts: 25 μL of the samples were used for immunoblot analysis, and 25 μL were used for nano flowcytometry analysis (nanoFCM).

### Immunoblot

Protein extracts were obtained after cell lysis in RIPA lysis buffer at 1:10 (Merck Millipore) supplemented with 0,1% SDS, a protease and phosphatase inhibitor cocktails (Roche). Protein quantification was performed by BCA assay (Thermo Scientific Pierce). Samples containing 30 µg of proteins were prepared with Laemmli loading buffer (Biorad) and DTT (Biorad), then heated at 70°C for 10 min. Protein were resolved by SDS-PAGE and allowed to migrate for 1 h 30 at 100 V. Proteins were then transferred to nitrocellulose membranes (Biorad) using liquid transfer at 100V during 1h. Blots were then blocked with 5% milk in PBS-Tween 20 0,1% (PBST) for 1 h at room temperature before overnight incubation at 4°C with specific primary antibodies (**Supplementary Table S3**). All primary antibodies were diluted in 5% w/v non-fat milk. After three 10 minute washes in PBST, membrane was incubated with appropriate HRP-conjugated secondary antibodies (1:10 000, Jackson Immuno Research) for 1 h at room temperature, followed by three washes and 5 min exposure in the dark to enhanced chemiluminescence signal (ECl, Biorad). A signal was acquired with a ChemiDoc XRS+ imaging system (Biorad) and blots were analyzed with Image Lab version 6.0.1 software.

### Image processing – Cell profiler

Image analyses were done using the Cell Profiler software (version 4.2.6) from the BROAD institute (the software is available at https://cellprofiler.org/).

### Segmentation process

- Step 1: MVBs in the images were segmented from the CD63 channel image using the IdentifyPrimaryObjects module

- Step 2: Nucleus in the images were segmented from the DAPI channel image using the IdentifyPrimaryObjects module

- Step 3: Cells in the images were segmented using the nucleus object (from step 2) and the cell mask channel image or a cytoplasmic protein staining using the IdentifySecondaryObjects module.

- Step 4: MVBs or other objects were link to their respective cell using the RelateObject module

- Step 5: To obtain intensity values the module MeasureObjectIntensity was used

- Step 6: To obtain area values the module MeasureObjectSizeShape was used

- Step 7: Measurements obtained from Step 5 and Step 6 were processed by the CalculateMath

- Step 8: All measurements were exported to excel file using the ExportTospReadsheat module

### Quantitative analysis of lipid species by shotgun lipidomics

After a 72-h siTFEB or siCtrl transfection, JMSU1 cells were harvested and resuspended in 200 µL of ammonium bicarbonate on ice. Samples were then snap-forzen in liquid N_2_ and stored at -80C until lipid analysis. Samples were spiked with 2.97 µL of a mixture of internal lipid standards containing 500 pmol of Chol-d6, 100 pmol of Chol-16:0-d7, 100 pmol of DAG 17:0-17:0, 50 pmol of TAG 17:0-17:0-17:0, 100 pmol of SM 18:1;2-12:0, 30 pmol of Cer 18:1;2-12:0, 30 pmol of GalCer 18:1;2-12:0, 50 pmol of LacCer 18:1;2-12:0, 300 pmol of PC 17:0-17:0, 50 pmol of PE 17:0-17:0, 50 pmol of PI 16:0-16:0, 50 pmol of PS 17:0-17:0, 30 pmol of PG 17:0-17:0, 30 pmol of PA 17:0-17:0, 40 pmol of Gb3 18:1:2-16:0-d9, 25 pmol of GM3 18:1;2-18:0-d5, 25 pmol of GM2 18:1;2-18:0-d9, 25 pmol of GM1 18:1;2-18:0-d5 and subjected to lipid extraction at 4 °C, as described elsewhere (Sampaio et al. 2011). Briefly, sample was extracted with 1 mL of chloroform-methanol (10:1) for 2 hours. The lower organic phase was collected, and the aqueous phase was re-extracted with 1 mL of chloroform-methanol (2:1) for 1 hour. The lower organic phase was collected and evaporated in a SpeedVac vacuum concentrator. Lipid extracts were dissolved in 100 μL of infusion mixture consisting of 7.5 mM ammonium acetate dissolved in propanol:chloroform:methanol [4:1:2 (vol/vol)].

Samples were analyzed by direct infusion in a QExactive Plus mass spectrometer (Thermo Fisher Scientific) equipped witha TriVersa NanoMate ion source (Advion Biosciences). 5 µL of sample were infused with gas pressure and voltage set to 1.25 psi and 0.95 kV, respectively. DAG, TAG and CE species were detected in the 10:1 extract, by positive ion mode FTMS as ammonium aducts by scanning m/z= 580–1000 Da, at R_m/z=200_=280 000 with lock mass activated at a common background (m/z=680.48022) for 30 seconds. Every scan is the average of 2 micro-scans, automatic gain control (AGC) was set to 1E6 and maximum ion injection time (IT) was set to 50ms

PC, PCO, Cer and GlcCer were detected as acetate adducts while PG, PE and PEO were detected as deprotonated adducts in the 10:1 extract, by negative ion mode FTMS, after polarity switch by scanning m/z= 420–1050 Da, at R_m/z=200_=280 000 with lock mass activated at a common background (m/z=529.46262) for 30 seconds. Every scan is the average of 2 micro-scans, automatic gain control (AGC) was set to 1E6 and maximum ion injection time (IT) was set to 50ms.

LacCer and Gb3 were detected as protonated ions and Gb4 was detected as ammoniated adduct in the 2:1 extract in positive ion mode FTMS by scanning m/z = 800–1,600 Da, at R_m/z=200_ = 280,000 with lock mass activated at a common background (m/z = 1,194.8179) for 30 s. GM1, GM2, andGM3 were detected as deprotonated ions in the 2:1 extract in negative ion mode after polarity switch in FTMS by scanning m/z = 1,100–1,650 Da, at R_m/z=200_= 280,000 with lock mass activated at a common background (m/z = 1,175.7768) for 30 s. Every scan is the average of two micro-scans, AGC was set to 1E6 and IT was set to 50 ms in both polarities.

PA, PI, PS, LPA, and LPS were detected as deprotonated ions in the 2:1 extract in negative ion mode in FTMS by scanning m/z = 400–1,100 Da, at R_m/z=200_ = 280,000 with lock mass activated at a common background (m/z = 529.4626) for 30 s. Every scan is the average of two micro-scans, AGC was set to 1E6 and IT was set to 50 ms. All data was acquired in centroid mode. All lipidomics data were analyzed with the lipid identification software, LipidXplorer (22272252 and 21247462). Tolerance for MS and identification was set to 2 ppm. Data post-processing and normalization to internal standards were done manually in Excel.

### Bodipy 493/503 staining

Lipid droplet content was quantified using the neutral lipid-specific probe Bodipy 493/503 (Invitrogen, Thermo Fisher Scientific). Briefly, TFEB-depleted cells were harvested, washed with 1X PBS, followed by incubation with Bodipy 493/503 at 1:1000 in 1X PBS for 20 min at room temperature. After staining is complete, the staining solution was replaced with prewarmed 1X PBS before analysis by flow cytometry on a BD FACSymphony A3 equipped with BD FACSDiva software (BD Biosciences). The data were analysed using FlowJo software (Tree Star).

Lipid droplets were also analyzed by fluorescence microscopy, after cell fixation in 4% PFA for 15 min at room temperature followed by washing and Bodipy incubation. Coverslips were then washed three times and mounted with Fluoromount-G (Thermo Fisher Scientific) supplemented with DAPI on glass slides. After drying overnight at 4°C, image acquisition was performed using a confocal Leica SP8.

### NR-Lyso probe

Endolysosomal membrane fluidity was analyzed using the acidic compartment-specific and polarity-sensitive probe NRlyso (Danylchuk et al. 2021). Briefly, cells were seeded on glass bottom dishes 35 mm (Ibidi). After treatments, cells were incubated with a mixture of 100 nM LysoTracker Blue (Invitrogen) and 50 nM NRLyso for 45 min at 37°C in serum-free and phenol red-free DMEM. Medium was then replaced with serum-free and phenol red-free medium supplemented with 20 mM HEPES (Thermo Fisher Scientific). As a positive control of membrane fluidity, cholesterol extraction was induced by incubating the cells with 10 mM methyl-β-cyclodextrin (mbCD, Sigma Aldrich) in RPMI for 30 min at 37°C.

The lipid order-dependent spectral shift of NRLyso was analyzed using a Leica SP8 confocal microscope with HC PL APO CS2 63×/1.40 OIL objective. For the first sequence, the excitation light was provided by a 488 nm laser, while the fluorescence was detected at two spectral ranges: 550−600 nm, on HyD detector 1 and 600−650 nm, on HyD detector 2, in sequential mode. For the second sequence, the LysoTracker blue probe was excited with a 405 nm laser and the signal detected at 415-480 nm on the HyD detector 1. The intensity ratio 550−600/600−650 was then analyzed on Fiji software.

### Laurdan probe

Total membrane fluidity was analyzed using the lipid fluorescent and polarity-sensitive probe Laurdan (Scheinpflug, Krylova, and Strahl 2017). Briefly, treated cells were harvested, washed with 1X PBS, followed by incubation with 200 nM Laurdan probe (Lumiprobe) in 1X PBS for 20 min at room temperature. After staining is complete, cells were washed two times and resuspended with prewarmed 1X PBS before analysis by flow cytometry on a LSR Fortessa (BD Biosciences). Laurdan is excited by a UV laser (355 nm) and the emitted fluorescence is recovered by the 488/10 nm and 450/40 nm filters. The median of fluorescence intensity (MFI) obtained at each emission wavelength is retrieved using the FlowJo® software and the MFI ratio at 490/440 nm is used to assess membrane fluidity.

For TFEB and HGS overexpression experiments, KU19-19 and JMSU1 cells were transiently transfected in a 12-well plate either with the pEGFP-N1-TFEB plasmid (#38119, Addgene), a kind gift from Shawn Ferguson, or the RFP-HGS plasmid (pCS2 HRS-RFP, a gift from Edward De Robertis (# 29685, Addgene) (Taelman et al. 2010), using the jetPEI reagent according to the manufacturer’s instructions (Polyplus). 48 h post-transfection, cells were harvested and MFI analysis was performed on GFP or RFP-positive cells.

### Exogenous lipid supplementation

Working solutions of oleic acid (OA, 18:1) (90260, Cayman Chemical) and palmitic acid (PA, 16:0) (10006627, Cayman Chemical) were diluted in 100% ethanol. Cells were then treated with these fatty acids diluted at a final concentration of 50 µM in complete RPMI medium for 24h, followed by subsequent analyses.

### Tumoroid formation and immunofluorescence staining

KU19-19 cells were seeded in bioengineered and soft hydrogel-containing micro-cavities of a 96-well plate in complete RPMI medium according to (Ouni et al. 2025). The next day, medium was replaced by a specific organoid culture medium previously described (Fujii et al. 2018; Cartry et al. 2023) allowing tumoroid growth. Following treatments, tumoroids were fixed in 4% PFA for 30 min at room temperature (RT) and three 10-min washes were performed. Next, fixed tumoroids were permeabilized in PBS, 0.2% Tritonx100 for 2 h at RT under agitation, followed by incubation in PBS 1X, 0.2% Tritonx100, 20% DMSO overnight at 37°C. The next day, autofluorescence quenching was performed by incubating tumoroids in sodium borohydride (1mg/mL) in PBS for 4 h at RT, washing 3 times for 30 min in PBS under agitation and leaving in PBS 1X overnight at RT. On day 3, tumoroids were incubated with a mix of primary antibodies (CD63 and ITGb5) diluted at 1:50 in PBS 1X, 1% FBS, 0.1% BSA, 0.2% Triton X100 for 24 h at 4°C under agitation. Following washing in 3% NaCl, 0.2% Tritonx100, PBS 1X (washing buffer) at 4°C under agitation for 4 h, tumoroids were incubated with secondary antibodies, goat anti-mouse Alexa Fluor 488 and anti-rabbit Alexa Fluor 568, diluted at 1:100 for 24 h at 4°C under agitation. Tumoroids were then washed with the washing buffer for 4 h, incubated with DAPI and Cell mask deep red actin stain (Invitrogen) at 1:1000 for 1 h at RT. After washing 2 times for 30 min in PBS under agitation, tumoroids were mounted with RapiClear 1.47 solution and keep at 4°C. The day of imaging, the plate was incubated at 37°C for 1 h before image acquisition on a spinning disk microscope (Gataca based on Nikon Eclipse Ti2).

### Data analysis from publicly available databases

Bioinformatic analysis of TFEB gene signature and HGS expression in bladder cancer was performed on patient samples from the PRISM study (Pradat et al. 2023).

### Gene Set Enrichment Analysis (GSEA)

The correlation between HGS expression and CLEAR network (genes list from Lachmann et al. 2010 was assessed using two independent approaches:

1. The 73 samples were divided into two groups based on HGS expression levels: low HGS expression (HGS Transcripts Per Kilobase Million [TPM] <40) and high HGS expression (HGS TPM > 40). The 40 TPM threshold was chosen because it splits the dataset into two nearly equal groups. The Log2 Fold Change (LFC) has been calculated for each gene between these two groups. Based on the LFC ranking, a GSEA for CLEAR network has been perform using the function fgsea() from the fgsea R package (preprint Korotkevich et al. 2021).
2. A GSEA for the CLEAR network has been performed for each sample independently, using the TPM-based gene expression ranking. The resulting Normalized Enrichment Score (NES) was then correlated to the natural logarithm of the normalized HGS expression. Normalized expression was calculated by dividing HGS TPM by the sample’s mean TPM. The correlation has been assessed using Pearson’s correlation coefficient and significance has been tested with a t-test for correlation, justified by the large sample size (n=73) and the apparent homoscedasticity.

### Statistics

GraphPad Prism 10.0 software was used for the graphical representation and statistical analysis of the data. Results were presented as mean ± s.d. of at least *N* = 3 independent experiments, unless stated otherwise in the figure legend. All the data were first tested for normality distribution to ensure appropriate statistical test usage. For all the figures, statistical significance was assessed at the 5% level and two-tailed unpaired statistical tests were performed. The results of the tests were displayed on the graphics and legends: non-significant (ns), * p<0.05, ** p<0.01, *** p<0.001.

## Acknowledgements

We thank the imaging and Cytometry core facility PFIC (SCR_028222) for assistance with microscopy. The imaging facility was supported by funding from the Paris-Saclay Cancer Cluster (PSCC) through an Agence Nationale de la Recherche (ANR) grant under the France 2030 program (ANR-22-BIOC-0001). We are grateful to the CurieCoreTech-Extracellular Vesicles Platform, Curie Institute, Paris, France for assistance with EVs isolation and analysis. KS received funding from Fondation Gusatve Roussy, ANR IntraTensionReg, La Ligue contre le Cancer and Inca PLBIO. JP received funding from Paris-Saclay. EO received funding from Paris Region Fellowship Programme, PRfP & Marie Skłodowska-Curie fellowship, AM received funding from Fondation de France.

## Author contributions

J.P. contributed to the design of the study, planned and conducted the experiments, prepared the figures and J.P. wrote original draft. A.M. planned and conducted the lipids fluidity and filipin experiments and prepared the figures. A.M. and E.O. performed 3D culture with microwell cavity assay. J.L.P and M.P. performed the lipidomic experiment and A.M. prepared the figures. H.L. performed the bioinformatics analyses on bladder cancer databases and J.P. assisted preparing the figures. N.E. and M.K. contributed to design of the study. G.B.B performed RNA sequencing on bladder cancer cell line. A.S.K provided NR-Lyso probe. K.S. contributed to the design of the study, coordinated the project, wrote original draft and reviewed it with the co-authors. The authors declare that they have no conflict of interest

---

**Supplementary Figure 1.**
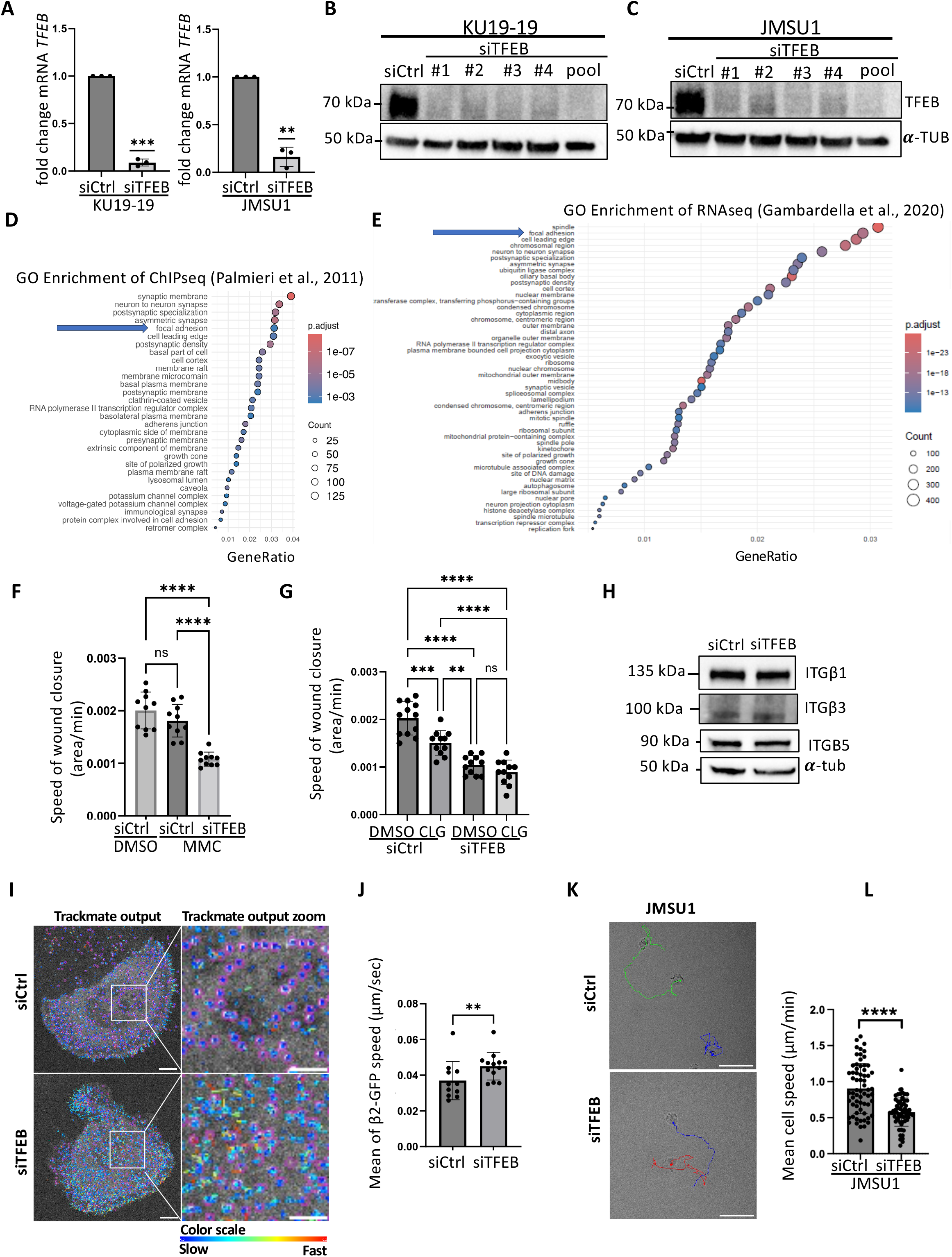
TFEB regulates cell migration and ITGβ5 trafficking. **(A)** RT-qPCR quantification of *TFEB* mRNA fold change in KU19-19 and JMSU1 cells treated with siControl (Ctrl) or siTFEB. Data represent mean ± SD from N=3; **p < 0.01, ***p < 0.001 (one-sample Student’s t test). **(B, C)** Immunoblot of TFEB in KU19-19 (B) and JMSU1 (C) cells treated with four individual siRNAs against TFEB (#1–#4) or a pooled siRNA (pool). α-Tubulin served as a loading control. **(D, E)** Gene Ontology analysis of TFEB targets from ChIP-seq (Palmeri et al., 2011) (D) and RNA-seq (Gambardella et al., 2020) (E). **(F)** Quantification of wound closure in KU19-19 cells transfected with siCtrl or siTFEB and treated with DMSO or mitomycin C (MMC, proliferation inhibitor). Wound area was calculated as |Tₙ–T₀|/T₀ in area/mins. Data represent mean ± SD from N=3; ns, not significant; ****p < 0.0001 (one-way ANOVA). **(G)** Quantification of wound closure in KU19-19 cells transfected with siCtrl or siTFEB and treated with DMSO or 10 μM cilengitide (CLG). Wound area was calculated as |Tₙ–T₀|/T₀ in area/mins. Data represent mean ± SD from N=3; ns, not significant; **p < 0.01, ***p < 0.001, ****p < 0.0001 (one-way ANOVA). **(H)** Immunoblot of ITGβ1, ITGβ3, and ITGβ5 in KU19-19 cells treated with siCtrl or siTFEB. α-Tubulin served as a loading control. **(I)** TIRF imaging of KU19-19 cells transfected with β2-GFP and siCtrl or siTFEB. Images were acquired every 5 s for 5 min. β2-GFP dynamics were analyzed using the TrackMate plugin in FIJI. Scale bars, 10 μm (overview) and 5 μm (zoom). Blue color represents slow tracks and red color represents fast tracks. **(J)** Quantification of β2-GFP speed in seconds/pix per image. Data represent mean ± SD from N=3; **p < 0.01 (two-tailed Student’s t test). **(K)** Single-cell manual tracking of JMSU1 cells transfected with siCtrl or siTFEB over 16 h. Scale bars, 100 μm. **(L)** Quantification of mean single-cell speed in μm/min. Data represent mean ± SD from N=3; ****p < 0.0001 (Student’s t test).

**Supplementary Figure 2.**
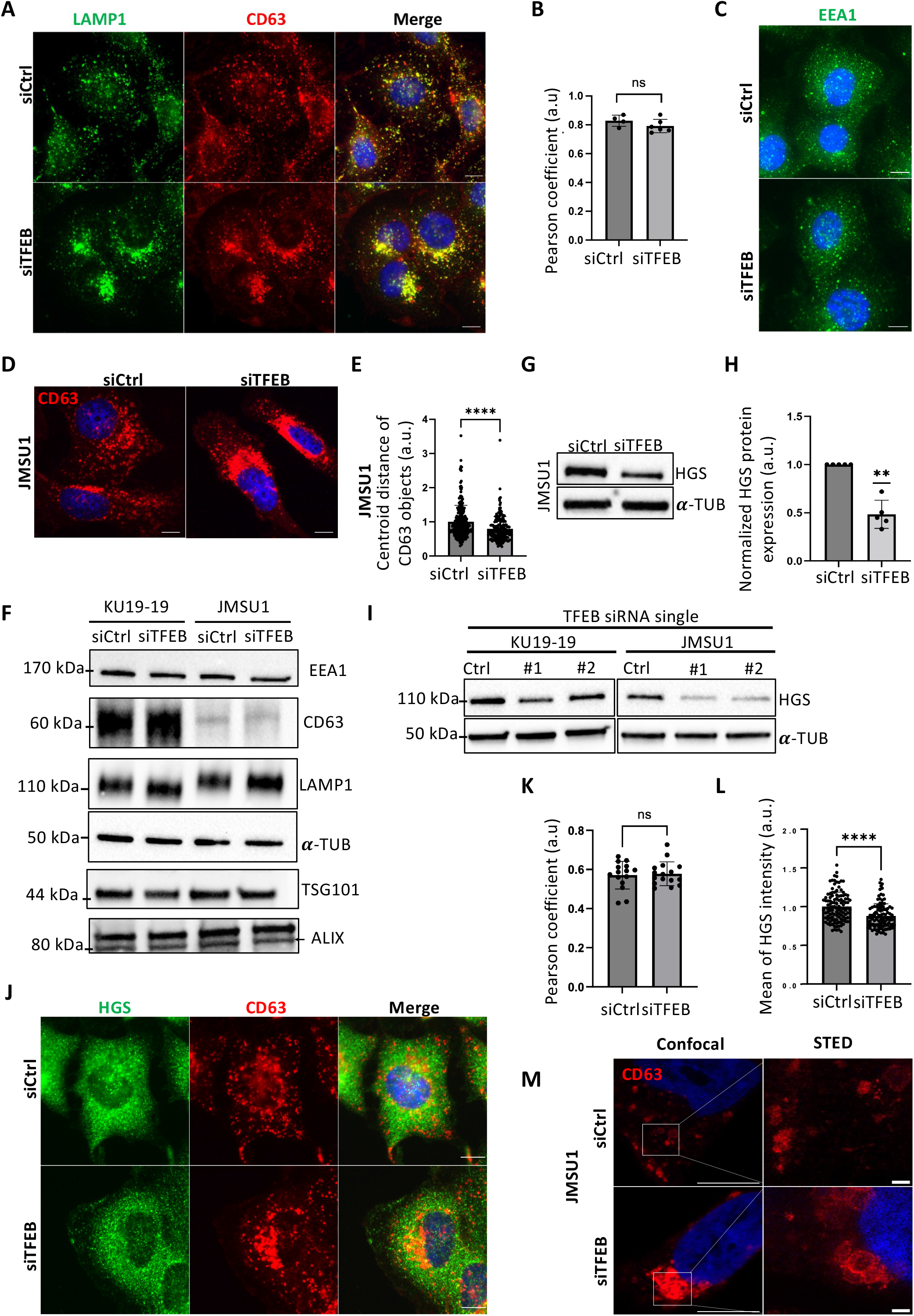
TFEB depletion promotes ITGβ5 accumulation in CD63-positive compartments and reduces HGS expression. **(A)** Immunofluorescence of CD63 (marker of MVBs) and LAMP1 (marker of lysosomes) in KU19-19 cells treated with siControl (Ctrl) or siTFEB. Nuclei were counterstained with DAPI. Scale bars, 10 μm. **(B)** Quantification of Pearson’s correlation coefficient for CD63 and LAMP1 colocalization in KU19-19 cells transfected with siCtrl or siTFEB. Data represent mean ± SD from N=1; ns not significant (Student’s t test). **(C)** Immunofluorescence of EEA1 (marker of early endosomes) in KU19-19 cells treated with siCtrl or siTFEB. Nuclei were stained with DAPI. Scale bars, 10 μm. **(D)** Immunofluorescence of CD63 in JMSU1 cells treated with siCtrl or siTFEB. Nuclei were stained with DAPI. Scale bars, 10 μm. **(E)** Quantification of the mean distance of CD63-positive structures to the cell centroid, normalized to cell area, in JMSU1 cells treated with siCtrl or siTFEB. Data represent mean ± SD from N=3; ****p < 0.0001 (Mann–Whitney test). **(F)** Immunoblot analysis of marker for early endosomes (EEA1), late endosomes/MVBs (CD63), lysosomes (LAMP1), and ESCRT components (ESCRT-I, TSG101; ESCRT-associated protein ALIX) in KU19-19 and JMSU1 cells treated with siCtrl or siTFEB. α-Tubulin served as a loading control. **(G)** Immunoblot of HGS in JMSU1 cells treated with siCtrl or siTFEB. α-Tubulin served as a loading control. **(H)** Densitometric quantification of HGS protein levels in JMSU1 cells treated with siCtrl or siHGS. Data represent mean ± SD from N=5; **p < 0.001 (one-sample Student’s t test). **(I)** Immunoblot of HGS in KU19-19 and JMSU1 cells treated with individual siRNAs targeting TFEB (#1 and #2; 50 nM, 72 h). **(J)** Immunofluorescence of CD63 (marker of MVBs) and HGS/Hrs (ESCRT-0) in KU19-19 cells treated with siControl (Ctrl) or siTFEB. Nuclei were counterstained with DAPI. Scale bars, 10 μm. **(K)** Quantification of Pearson’s correlation coefficient for CD63 and HGS/Hrs colocalization in KU19-19 cells transfected with siCtrl or siTFEB (arbitrary units, a.u.). Data represent mean ± SD from N=2; ns not significant (Student’s t test). **(L)** Quantification of mean HGS intensity normalized to cell area in KU19-19 cells treated with siControl (Ctrl) or siTFEB. Data represent mean ± SD from N=2; ****p < 0.0001 (Mann–Whitney test). **(M)** Images from stimulated emission depletion (STED) microscopy of CD63 in JMSU1 cells treated with siCtrl or siTFEB. Nuclei were stained with DAPI. Scale bars, 10 μm (overview) and 1 μm (inset).

**Supplementary Figure 3.**
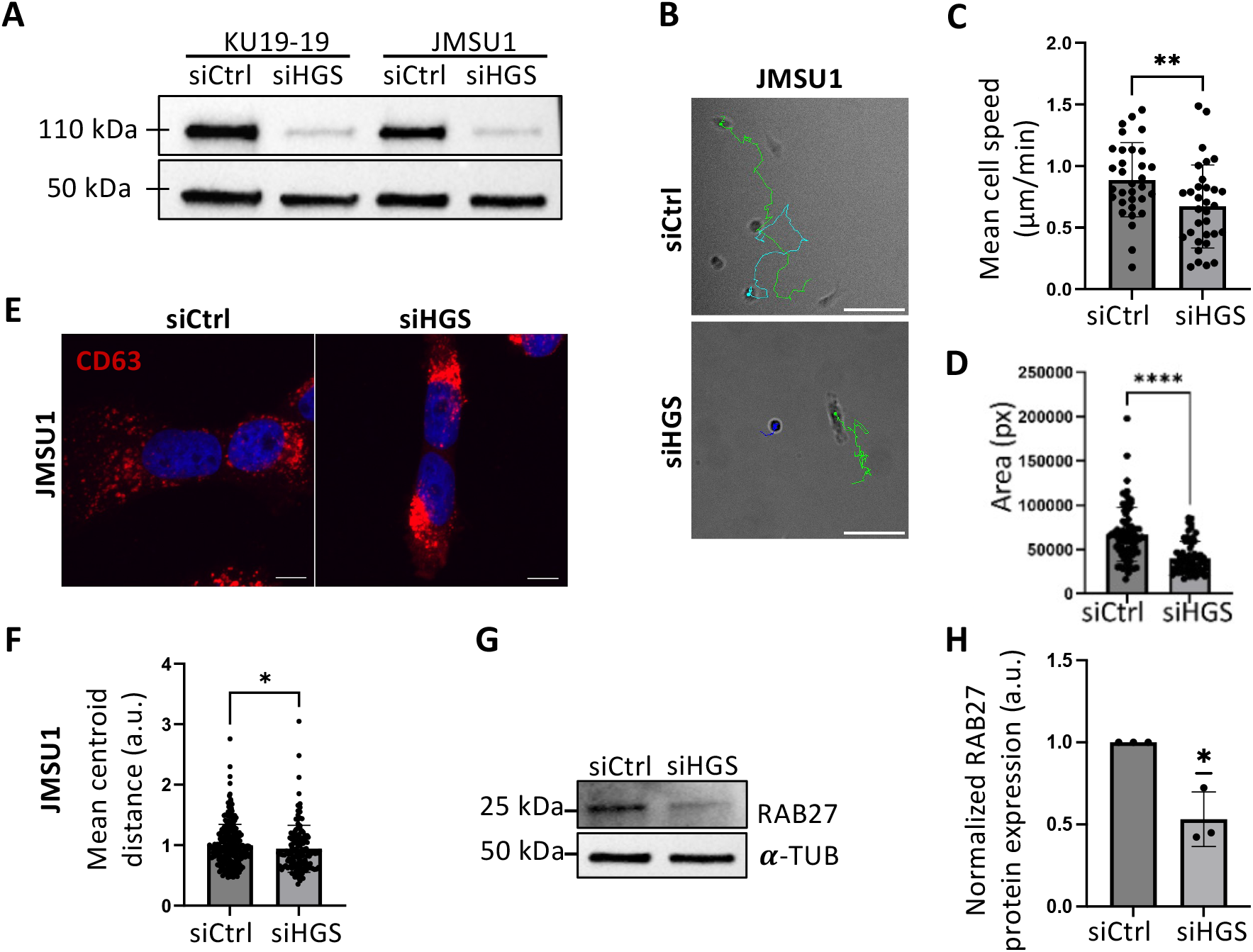
ESCRT-0 HGS/Hrs regulates ITGβ5 trafficking downstream of TFEB. (A) Immunoblot analysis of HGS in KU19-19 and JMSU1 cells transfected with siControl (Ctrl) or siHGS. α-Tubulin served as a loading control. (B) Representative single-cell manual tracking analysis of JMSU1 cells transfected with siCtrl or siHGS. (C) Quantification of mean single-cell migration speed in μm/min. Data represent mean ± SD of N=3; **p < 0.01 (Student’s t test). (D) Quantification of individual cell area in pixels (px) in JMSU1 cells treated with siCtrl or siHGS. Data represent mean ± SD of N=3; ****p < 0.0001 (Mann–Whitney test). (E) Immunofluorescence staining of CD63 (marker of multivesicular bodies) in JMSU1 cells treated with siCtrl or siHGS. Nuclei were counterstained with DAPI. Scale bars, 10 μm. (F) Quantification of the mean distance of CD63-positive objects to the cell centroid, normalized to cell area in arbitrary units (a.u.), in JMSU1 cells treated with siCtrl or siHGS. Data represent mean ± SD of N=3; *p < 0.05 (Mann–Whitney test). (G) Immunoblot analysis of Rab27 in JMSU1 cells treated with siCtrl or siHGS. α-Tubulin served as a loading control. (H) Densitometric quantification of RAB27 protein levels from immunoblots in JMSU1 cells treated with siCtrl or siHGS in arbitrary units (a.u.). Data represent mean ± SD of N=3; *p < 0.05 (One sample Student’s t test).

**Supplementary Figure 4.**
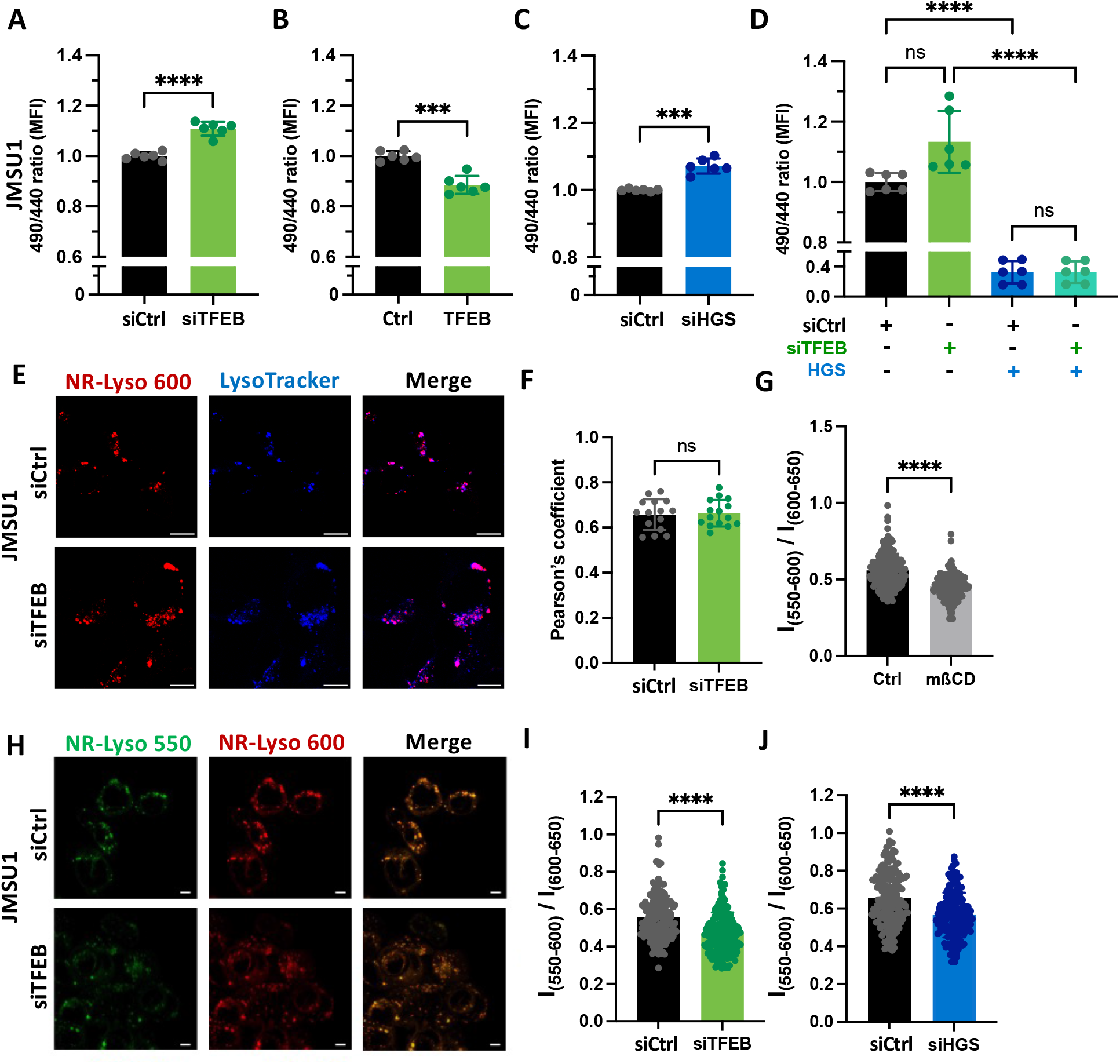
TFEB and HGS/Hrs regulate endosomal membrane fluidity. (A) Ratiometric measurement of membrane fluidity by flow cytometry using the polarity-sensitive Laurdan dye in JMSU1 cells 72 h post-transfection with siControl (Ctrl) or siTFEB. Data represent mean ± SD of N=3; ****p < 0.0001 (unpaired Student’s t test with Welch’s correction). (B) Ratiometric measurement of membrane fluidity by flow cytometry using the polarity-sensitive Laurdan dye in JMSU1 cells 24 h post-transfection with the EGFP vector (Ctrl) or pEGFP-TFEB plasmid (TFEB). Data represent mean ± SD of N=3; ***p < 0.001 (unpaired Student’s t test with Welch’s correction). (C) Ratiometric measurement of membrane fluidity by flow cytometry using Laurdan dye in JMSU1 cells 48 h post-transfection with siHGS or siCtrl. Data represent mean ± SD of N=3; ***p < 0.001 (unpaired Student’s t test with Welch’s correction). (D) Ratiometric measurement of membrane fluidity by flow cytometry using Laurdan dye in JMSU1 cells 72 h post-transfection with siTFEB or siCtrl and 24 h post-transfection with the pCS2-HRS-EGFP plasmid (HGS) or EGFP vector (Ctrl). Data represent mean ± SD of N=3; n.s. not significant; ****p < 0.0001 (unpaired Student’s t test with Welch’s correction). (E) Fluorescence staining of NR-Lyso (red) and LysoTracker (blue) in JMSU1 cells treated with siCtrl or siTFEB. Scale bars, 10 μm. **(F)** Quantification of Pearson’s correlation coefficient for NR-Lyso and LysoTracker colocalization in JMSU1 cells transfected with siCtrl or siTFEB (arbitrary units, a.u.). Data represent mean ± SD from N=3; ns not significant (Student’s t test). (G) Quantification of the green/red fluorescence intensity ratio (I₅₅₀–₆₀₀/I₆₀₀–₆₅₀) of the NR-Lyso probe in JMSU1 cells treated for 30 min with 10 mM methyl-β-cyclodextrin (mβCD). Data represent mean ± SD of N=3; ****p < 0.0001 (Mann– Whitney U test). (H) Representative ratiometric confocal images of JMSU1 cells stained with NR-Lyso. Scale bars, 10 μm. (I) Quantification of the green/red fluorescence intensity ratio (I₅₅₀–₆₀₀/I₆₀₀–₆₅₀) of the NR-Lyso probe in JMSU1 cells transfected with siCtrl or siTFEB. Data represent mean ± SD of three independent experiments; ****p < 0.0001 (Mann–Whitney U test). (J) Quantification of the green/red fluorescence intensity ratio (I₅₅₀–₆₀₀/I₆₀₀–₆₅₀) of the NR-Lyso probe in JMSU1 cells transfected with siCtrl or siHGS. Data represent mean ± SD of N=3; ****p < 0.0001 (Mann–Whitney U test).

**Supplementary Figure 5.**
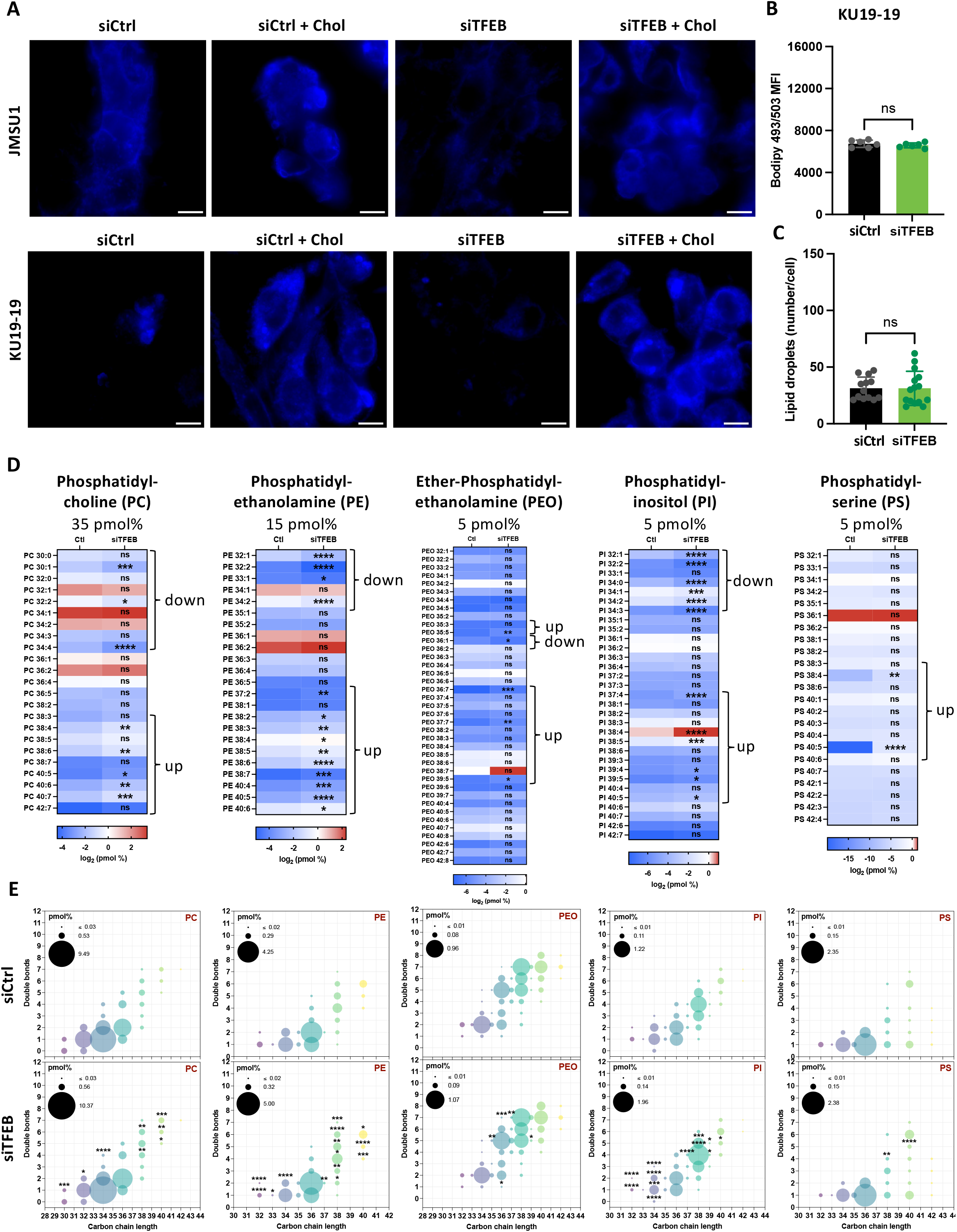
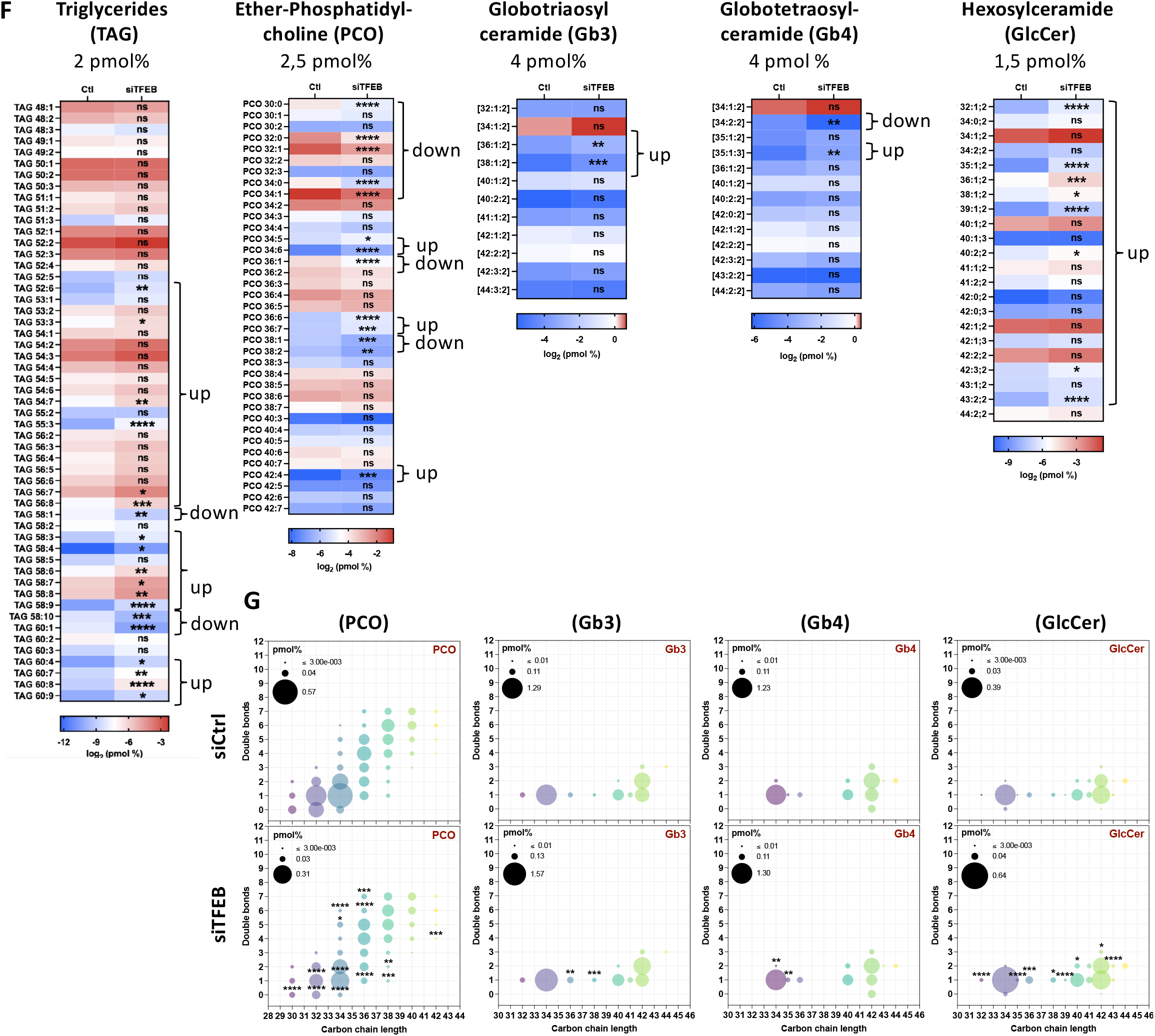
TFEB regulates lipid composition in JMSU1 cells. (A) Representative confocal images of Filipin fluorescence in control and TFEB-depleted JMSU1 and KU19-19 cells 72 h post-transfection. Cholesterol was added for 3h when indicated. Scale bars, 10 μm. (B) Flow cytometry analysis of lipid droplet content in control and TFEB-depleted KU19-19 cells stained with Bodipy 493/503 (green) 72 h post-transfection. Data represent mean ± SD of N=3; ns, not significant (unpaired Student’s t test with Welch’s correction). (C) Quantification of lipid droplet number per cell in KU19-19. Data represent mean ± SD of N=3; ns, not significant (Mann–Whitney U test). (D) Heatmaps showing the relative changes of abundant (>5 pmol%) phospholipids including phosphatidylcholine (PC), phosphatidylethanolamine (PEO), phosphatidylinositol (PI) and phosphatidylserine (PS) in control and TFEB-depleted JMSU1 cells. Red indicates increased levels, and blue indicates decreased levels. Data are expressed as log₂-transformed lipid quantities (pmol %). Significance was determined by an unpaired Student’s t test with Welch’s correction; ns, not significant; *p < 0.05; **p < 0.01; ***p < 0.001; ****p < 0.0001. (E) Bubble plot of abundant (>5 pmol%) phospholipids including phosphatidylcholine (PC), phosphatidylethanolamine (PEO), phosphatidylinositol (PI) and phosphatidylserine (PS) in control (upper panel) and TFEB-depleted (lower panel) JMSU1 cells. Each circle represents a phospholipid species defined by its number of carbon atoms and double bonds. Circle size indicates the average quantity of each species, and fill color represents total carbon content. Significance was determined by two-way ANOVA with Šídák’s multiple comparisons test; *p < 0.05; **p < 0.01; ***p < 0.001; ****p < 0.0001. (F) Heatmaps showing the relative changes of different lipid species (>1,5 pmol%), namely, triglycerides (TAG), ether-phosphatidylcholine (PCO) and the sphingolipids globotriaosylceramide (Gb3), globotetraosylceramide (Gb4) and hexosylceramide (GlcCer) in control and TFEB-depleted JMSU1 cells. Red indicates increased levels, and blue indicates decreased levels. Data are expressed as log₂-transformed lipid quantities (pmol %). Significance was determined by an unpaired t test with Welch’s correction; ns, not significant; *p < 0.05; **p < 0.01; ***p < 0.001; ****p < 0.0001. (H) Bubble plot of different lipid species (>1,5 pmol%), namely, ether-phosphatidylcholine (PCO), globotriaosylceramide (Gb3), globotetraosylceramide (Gb4) and hexosylceramide (GlcCer) in control (upper panel) and TFEB-depleted (lower panel) JMSU1 cells. Each circle represents a phospholipid species defined by its number of carbon atoms and double bonds. Circle size indicates the average quantity of each species, and fill color represents total carbon content. Significance was determined by two-way ANOVA with Šídák’s multiple comparisons test; *p < 0.05; **p < 0.01; ***p < 0.001; ****p < 0.0001.

**Supplementary Figure 6.**
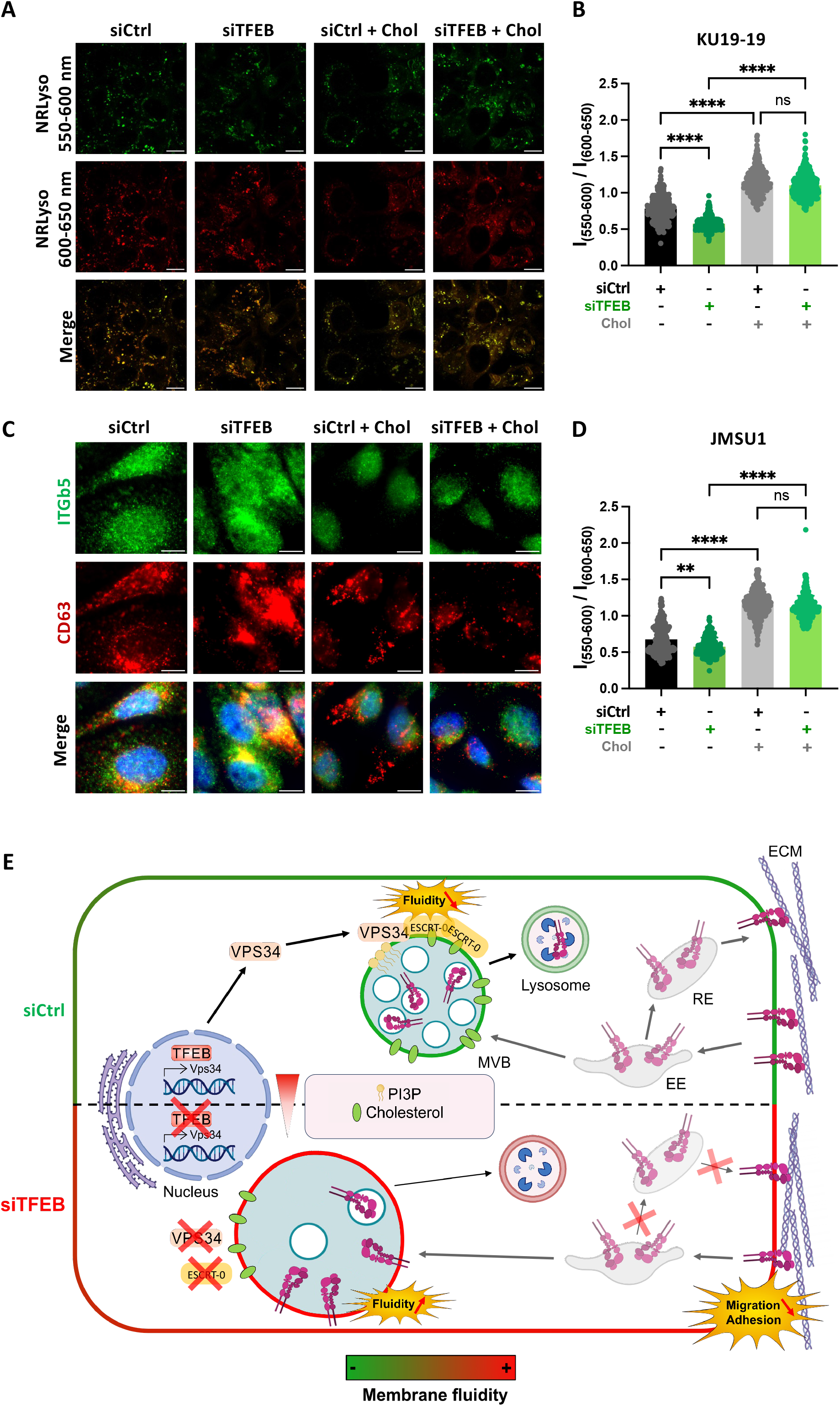
Fatty acid–regulated endosomal membrane fluidity controls MVB positioning and ITGβ5 trafficking. (A) Representative ratiometric confocal images of NR-Lyso-stained KU19-19 cells transfected with siCtrl or siTFEB and treated for 24 h with ethanol 100% (Ctrl) or cholesterol (Chol). (B) Quantification of the green/red fluorescence intensity ratio (I₅₅₀–₆₀₀ / I₆₀₀–₆₅₀) in KU19-19 cells transfected with siCtrl or siTFEB and treated for 24 h with ethanol 100% (Ctrl) or cholesterol (Chol). Data represent mean ± SD of N=3; ns, not significant, ****p < 0.0001 (Kruskal–Wallis test followed by Dunn’s multiple comparisons). (C) Confocal images of KU19-19 cells transfected with siCtrl or siTFEB and treated for 24 h with ethanol 100% (Ctrl) or cholesterol (Chol) and immunostained for CD63 (red) and ITGβ5 (green). Nuclei were counterstained with DAPI. Scale bars, 10 μm.Scale bars, 10 μm. (D) Quantification of the green/red fluorescence intensity ratio (I₅₅₀–₆₀₀ / I₆₀₀–₆₅₀) in JMSU1 cells transfected with siCtrl or siTFEB and treated for 24 h with ethanol 100% (Ctrl) or cholesterol (Chol). Data represent mean ± SD of N=3; ns, not significant, ****p < 0.0001 (Kruskal–Wallis test followed by Dunn’s multiple comparisons).

## Bibliography

1. Alonso-Bivou, Mariano, Albert Pol, and Harriet P. Lo. 2025. “Moving the Fat: Emerging Roles of Rab GTPases in the Regulation of Lipid Droplet Contact Sites.” Current Opinion in Cell Biology 93: 102466. doi:10.1016/j.ceb.2025.102466.

2. Bakouny, Ziad, Ananthan Sadagopan, Praful Ravi, Nebiyou Y. Metaferia, Jiao Li, Shatha AbuHammad, Stephen Tang, et al. 2022. “Integrative Clinical and Molecular Characterization of Translocation Renal Cell Carcinoma.” Cell Reports 38(1): 110190. doi:10.1016/j.celrep.2021.110190.

3. Ballabio, Andrea. 2016. “The Awesome Lysosome.” EMBO molecular medicine 8(2): 73–76. doi:10.15252/emmm.201505966.

4. Baschieri, Francesco, Stéphane Dayot, Nadia Elkhatib, Nathalie Ly, Anahi Capmany, Kristine Schauer, Timo Betz, et al. 2018. “Frustrated Endocytosis Controls Contractility-Independent Mechanotransduction at Clathrin-Coated Structures.” Nature Communications 9(1): 3825. doi:10.1038/s41467-018-06367-y.

5. Beckmann, H., L. K. Su, and T. Kadesch. 1990. “TFE3: A Helix-Loop-Helix Protein That Activates Transcription through the Immunoglobulin Enhancer muE3 Motif.” Genes & Development 4(2): 167–79. doi:10.1101/gad.4.2.167.

6. Bergen, Janice, Martina Karasova, Andrea Bileck, Marc Pignitter, Doris Marko, Christopher Gerner, and Giorgia Del Favero. 2023. “Exposure to Dietary Fatty Acids Oleic and Palmitic Acid Alters Structure and Mechanotransduction of Intestinal Cells in Vitro.” Archives of Toxicology 97(6): 1659–75. doi:10.1007/s00204-023-03495-3.

7. Calcagnì, Alessia, Lotte Kors, Eric Verschuren, Rossella De Cegli, Nicolina Zampelli, Edoardo Nusco, Stefano Confalonieri, et al. 2016. “Modelling TFE Renal Cell Carcinoma in Mice Reveals a Critical Role of WNT Signaling.” eLife 5: e17047. doi:10.7554/eLife.17047.

8. Calderwood, David A., Iain D. Campbell, and David R. Critchley. 2013. “Talins and Kindlins: Partners in Integrin-Mediated Adhesion.” Nature Reviews. Molecular Cell Biology 14(8): 503–17. doi:10.1038/nrm3624.

9. Carpentier, Maxime, Mohyeddine Omrane, Rola Shaaban, Jennica Träger, Naïma El Khallouki, Soazig Le Lay, Julie Patat, et al. 2025. “Seipin Governs Caveolin-1 Trafficking through Modulating Sphingolipid-Glycerolipid Balance.” Cell Reports 44(10): 116320. doi:10.1016/j.celrep.2025.116320.

10. Carr, C S, and P A Sharp. 1990. “A Helix-Loop-Helix Protein Related to the Immunoglobulin E Box-Binding Proteins.” Molecular and Cellular Biology 10(8): 4384–88. doi:10.1128/mcb.10.8.4384.

11. Cartry, Jérôme, Sabrina Bedja, Alice Boilève, Jacques R. R. Mathieu, Emilie Gontran, Maxime Annereau, Bastien Job, et al. 2023. “Implementing Patient Derived Organoids in Functional Precision Medicine for Patients with Advanced Colorectal Cancer.” Journal of Experimental & Clinical Cancer Research : CR 42: 281. doi:10.1186/s13046-023-02853-4.

12. Chastney, Megan R., Jasmin Kaivola, Veli-Matti Leppänen, and Johanna Ivaska. 2025. “The Role and Regulation of Integrins in Cell Migration and Invasion.” Nature Reviews. Molecular Cell Biology 26(2): 147–67. doi:10.1038/s41580-024-00777-1.

13. Chen, Lupeng, Yue Liu, Junzhi Zhang, Tongxing Song, Jian Wu, and Zhuqing Ren. 2025. “AMPK Regulates ARF1 Localization to Membrane Contact Sites to Facilitate Fatty Acid Transfer between Lipid Droplets and Mitochondria.” Cell Death & Disease 16(1): 623. doi:10.1038/s41419-025-07957-7.

14. Covino, Roberto, Gerhard Hummer, and Robert Ernst. 2018. “Integrated Functions of Membrane Property Sensors and a Hidden Side of the Unfolded Protein Response.” Molecular Cell 71(3): 458–67. doi:10.1016/j.molcel.2018.07.019.

15. Da Graça, Juliane, Cédric Delevoye, and Etienne Morel. 2025. “Morphodynamical Adaptation of the Endolysosomal System to Stress.” The FEBS Journal 292(2): 248–60. doi:10.1111/febs.17154.

16. Danylchuk, Dmytro I., Pierre-Henri Jouard, and Andrey S. Klymchenko. 2021. “Targeted Solvatochromic Fluorescent Probes for Imaging Lipid Order in Organelles under Oxidative and Mechanical Stress.” Journal of the American Chemical Society 143(2): 912–24. doi:10.1021/jacs.0c10972.

17. Darwich, Zeinab, Andrey S. Klymchenko, Denis Dujardin, and Yves Mély. 2014. “Imaging Lipid Order Changes in Endosome Membranes of Live Cells by Using a Nile Red-Based Membrane Probe.” RSC Advances 4(17): 8481–88. doi:10.1039/C3RA47181K.

18. De Santis, Augusta, Sara La Manna, Irene Russo Krauss, Anna Maria Malfitano, Ettore Novellino, Luca Federici, Antonella De Cola, et al. 2018. “Nucleophosmin-1 Regions Associated with Acute Myeloid Leukemia Interact Differently with Lipid Membranes.” Biochimica Et Biophysica Acta. General Subjects 1862(4): 967–78. doi:10.1016/j.bbagen.2018.01.005.

19. Di Malta, Chiara, Angela Zampelli, Letizia Granieri, Claudia Vilardo, Rossella De Cegli, Laura Cinque, Edoardo Nusco, et al. 2023. “TFEB and TFE3 Drive Kidney Cystogenesis and Tumorigenesis.” EMBO Molecular Medicine 15(5): e16877. doi:10.15252/emmm.202216877.

20. Diamant, Ido, Daniel J B Clarke, John Erol Evangelista, Nathania Lingam, and Avi Ma’ayan. 2025. “Harmonizome 3.0: Integrated Knowledge about Genes and Proteins from Diverse Multi-Omics Resources.” Nucleic Acids Research 53(D1): D1016–28. doi:10.1093/nar/gkae1080.

21. Doronzo, Gabriella, Elena Astanina, Davide Corà, Giulia Chiabotto, Valentina Comunanza, Alessio Noghero, Francesco Neri, et al. 2019. “TFEB Controls Vascular Development by Regulating the Proliferation of Endothelial Cells.” The EMBO Journal 38(3): e98250. doi:10.15252/embj.201798250.

22. Edgar, James R., Emily R. Eden, and Clare E. Futter. 2014. “Hrs- and CD63-Dependent Competing Mechanisms Make Different Sized Endosomal Intraluminal Vesicles.” *Traffic (Copenhagen*, Denmark) 15(2): 197–211. doi:10.1111/tra.12139.

23. Ehringer, W., D. Belcher, S. R. Wassall, and W. Stillwell. 1990. “A Comparison of the Effects of Linolenic (18:3 Omega 3) and Docosahexaenoic (22:6 Omega 3) Acids on Phospholipid Bilayers.” Chemistry and Physics of Lipids 54(2): 79–88. doi:10.1016/0009-3084(90)90063-w.

24. Elkhatib, Nadia, Enzo Bresteau, Francesco Baschieri, Alba López Rioja, Guillaume van Niel, Stéphane Vassilopoulos, and Guillaume Montagnac. 2017. “Tubular Clathrin/AP-2 Lattices Pinch Collagen Fibers to Support 3D Cell Migration.” Science 356(6343): eaal4713. doi:10.1126/science.aal4713.

25. Erlich, Avigail T., Diane M. Brownlee, Kaitlyn Beyfuss, and David A. Hood. 2018. “Exercise Induces TFEB Expression and Activity in Skeletal Muscle in a PGC-1α-Dependent Manner.” American Journal of Physiology. Cell Physiology 314(1): C62–72. doi:10.1152/ajpcell.00162.2017.

26. Evans, Trent D., Xiangyu Zhang, Se-Jin Jeong, Anyuan He, Eric Song, Somashubhra Bhattacharya, Karyn B. Holloway, Irfan J. Lodhi, and Babak Razani. 2019. “TFEB Drives PGC-1α Expression in Adipocytes to Protect against Diet-Induced Metabolic Dysfunction.” Science Signaling 12(606): eaau2281. doi:10.1126/scisignal.aau2281.

27. Fujii, Masayuki, Mami Matano, Kohta Toshimitsu, Ai Takano, Yohei Mikami, Shingo Nishikori, Shinya Sugimoto, and Toshiro Sato. 2018. “Human Intestinal Organoids Maintain Self-Renewal Capacity and Cellular Diversity in Niche-Inspired Culture Condition.” Cell Stem Cell 23(6): 787–793.e6. doi:10.1016/j.stem.2018.11.016.

28. Gambardella, Gennaro, Leopoldo Staiano, Maria Nicoletta Moretti, Rossella De Cegli, Luca Fagnocchi, Giuseppe Di Tullio, Sara Polletti, et al. 2020. “GADD34 Is a Modulator of Autophagy during Starvation.” Science Advances 6(39): eabb0205. doi:10.1126/sciadv.abb0205.

29. Gaullier, Jean-Michel, Anne Simonsen, Antonello D’Arrigo, Bjørn Bremnes, Harald Stenmark, and Rein Aasland. 1998. “FYVE Fingers Bind PtdIns(3)P.” Nature 394(6692): 432–33. doi:10.1038/28767.

30. Ge, Yifan, Jiayun Gao, Rainer Jordan, and Christoph A. Naumann. 2018. “Changes in Cholesterol Level Alter Integrin Sequestration in Raft-Mimicking Lipid Mixtures.” Biophysical Journal 114(1): 158–67. doi:10.1016/j.bpj.2017.11.005.

31. Giatromanolaki, Alexandra, Dimitra Kalamida, Efthimios Sivridis, Ilias V. Karagounis, Kevin C. Gatter, Adrian L. Harris, and Michael I. Koukourakis. 2015. “Increased Expression of Transcription Factor EB (TFEB) Is Associated with Autophagy, Migratory Phenotype and Poor Prognosis in Non-Small Cell Lung Cancer.” *Lung Cancer (Amsterdam*, Netherlands) 90(1): 98–105. doi:10.1016/j.lungcan.2015.07.008.

32. Gilleron, Jerome, and Anja Zeigerer. 2023. “Endosomal Trafficking in Metabolic Homeostasis and Diseases.” Nature Reviews. Endocrinology 19(1): 28–45. doi:10.1038/s41574-022-00737-9.

33. Gopaldass, Navin, Kai-En Chen, Brett Collins, and Andreas Mayer. 2024. “Assembly and Fission of Tubular Carriers Mediating Protein Sorting in Endosomes.” Nature Reviews. Molecular Cell Biology 25(10): 765–83. doi:10.1038/s41580-024-00746-8.

34. Hakala, Markku, Satish Babu Moparthi, Iva Ganeva, Mehmet Gül, César Bernat-Silvestre, Carlos Marcuello, Javier Espadas, et al. 2026. “Two-Dimensional HRS Condensates Drive the Assembly of Flat Clathrin Lattices on Endosomes.” Nature Communications. doi:10.1038/s41467-026-73132-x.

35. Harayama, Takeshi, and Bruno Antonny. 2023. “Beyond Fluidity: The Role of Lipid Unsaturation in Membrane Function.” Cold Spring Harbor Perspectives in Biology 15(7): a041409. doi:10.1101/cshperspect.a041409.

36. Hertwig, Paula. 1942. “Neue Mutationen und Koppelungsgruppen bei der Hausmaus.” Zeitschrift für Induktive Abstammungs- und Vererbungslehre 80(1): 220–46. doi:10.1007/BF01741984.

37. Huan, Chongmin, Matthew L Kelly, Ryan Steele, Iuliana Shapira, Susan R S Gottesman, and Christopher A J Roman. 2006. “Transcription Factors TFE3 and TFEB Are Critical for CD40 Ligand Expression and Thymus-Dependent Humoral Immunity.” Nature immunology 7(10): 1082–91. doi:10.1038/ni1378.

38. Huan, Chongmin, Deepa Sashital, Tiruneh Hailemariam, Matthew L. Kelly, and Christopher A. J. Roman. 2005. “Renal Carcinoma-Associated Transcription Factors TFE3 and TFEB Are Leukemia Inhibitory Factor-Responsive Transcription Activators of E-Cadherin.” The Journal of Biological Chemistry 280(34): 30225–35. doi:10.1074/jbc.M502380200.

39. Huotari, Jatta, and Ari Helenius. 2011. “Endosome Maturation.” The EMBO Journal 30(17): 3481–3500. doi:10.1038/emboj.2011.286.

40. Jain, Aakriti, and Roberto Zoncu. 2026. “Lysosomes as Hubs of Metabolic Sensing and Cellular Homeostasis.” Molecular Cell 86(3): 533–52. doi:10.1016/j.molcel.2026.01.011.

41. Jian, Ye, Shunling Yuan, Jialun Yang, Yong Lei, Xuan Li, and Wenfeng Liu. 2022. “Aerobic Exercise Alleviates Abnormal Autophagy in Brain Cells of APP/PS1 Mice by Upregulating AdipoR1 Levels.” International Journal of Molecular Sciences 23(17): 9921. doi:10.3390/ijms23179921.

42. Kim, Soyoung, Gahyeon Song, Taebok Lee, Minseong Kim, Jeongrae Kim, Hyeryun Kwon, Jiyoung Kim, et al. 2021. “PARsylated Transcription Factor EB (TFEB) Regulates the Expression of a Subset of Wnt Target Genes by Forming a Complex with β-Catenin-TCF/LEF1.” Cell Death and Differentiation 28(9): 2555–70. doi:10.1038/s41418-021-00770-7.

43. Klymchenko, Andrey S. 2017. “Solvatochromic and Fluorogenic Dyes as Environment-Sensitive Probes: Design and Biological Applications.” Accounts of Chemical Research 50(2): 366–75. doi:10.1021/acs.accounts.6b00517.

44. Klymchenko, Andrey S. 2023. “Fluorescent Probes for Lipid Membranes: From the Cell Surface to Organelles.” Accounts of Chemical Research 56(1): 1–12. doi:10.1021/acs.accounts.2c00586.

45. Kobayashi, T., M. H. Beuchat, M. Lindsay, S. Frias, R. D. Palmiter, H. Sakuraba, R. G. Parton, and J. Gruenberg. 1999. “Late Endosomal Membranes Rich in Lysobisphosphatidic Acid Regulate Cholesterol Transport.” Nature Cell Biology 1(2): 113–18. doi:10.1038/10084.

46. Korotkevich, Gennady, Vladimir Sukhov, Nikolay Budin, Boris Shpak, Maxim N. Artyomov, and Alexey Sergushichev. 2021. “Fast Gene Set Enrichment Analysis.” : 060012. doi:10.1101/060012.

47. Kucherak, Oleksandr A., Sule Oncul, Zeinab Darwich, Dmytro A. Yushchenko, Youri Arntz, Pascal Didier, Yves Mély, and Andrey S. Klymchenko. 2010. “Switchable Nile Red-Based Probe for Cholesterol and Lipid Order at the Outer Leaflet of Biomembranes.” Journal of the American Chemical Society 132(13): 4907–16. doi:10.1021/ja100351w.

48. Lachmann, Alexander, Huilei Xu, Jayanth Krishnan, Seth I. Berger, Amin R. Mazloom, and Avi Ma’ayan. 2010. “ChEA: Transcription Factor Regulation Inferred from Integrating Genome-Wide ChIP-X Experiments.” Bioinformatics 26(19): 2438–44. doi:10.1093/bioinformatics/btq466.

49. Lietha, Daniel, and Tina Izard. 2020. “Roles of Membrane Domains in Integrin-Mediated Cell Adhesion.” International Journal of Molecular Sciences 21(15): 5531. doi:10.3390/ijms21155531.

50. Lingwood, Daniel, and Kai Simons. 2010. “Lipid Rafts As a Membrane-Organizing Principle.” Science 327(5961): 46–50. doi:10.1126/science.1174621.

51. Lobert, Viola Hélène, Andreas Brech, Nina Marie Pedersen, Jørgen Wesche, Angela Oppelt, Lene Malerød, and Harald Stenmark. 2010. “Ubiquitination of Α5β1 Integrin Controls Fibroblast Migration through Lysosomal Degradation of Fibronectin-Integrin Complexes.” Developmental Cell 19(1): 148–59. doi:10.1016/j.devcel.2010.06.010.

52. Lobert, Viola Hélène, and Harald Stenmark. 2012. “The ESCRT Machinery Mediates Polarization of Fibroblasts through Regulation of Myosin Light Chain.” Journal of Cell Science 125(1): 29–36. doi:10.1242/jcs.088310.

53. MacDonald, Ewan, Louise Brown, Arnaud Selvais, Han Liu, Thomas Waring, Daniel Newman, Jessica Bithell, et al. 2018. “HRS–WASH Axis Governs Actin-Mediated Endosomal Recycling and Cell Invasion.” Journal of Cell Biology 217(7): 2549–64. doi:10.1083/jcb.201710051.

54. Manni, Marco M., Jesús Sot, Enara Arretxe, Rubén Gil-Redondo, Juan M. Falcón-Pérez, David Balgoma, Cristina Alonso, Félix M. Goñi, and Alicia Alonso. 2018. “The Fatty Acids of Sphingomyelins and Ceramides in Mammalian Tissues and Cultured Cells: Biophysical and Physiological Implications.” Chemistry and Physics of Lipids 217: 29–34. doi:10.1016/j.chemphyslip.2018.09.010.

55. Mao, Xiaodan, Huifang Lei, Tianjin Yi, Pingping Su, Shuting Tang, Yao Tong, Binhua Dong, et al. 2022. “Lipid Reprogramming Induced by the TFEB-ERRα Axis Enhanced Membrane Fluidity to Promote EC Progression.” Journal of Experimental & Clinical Cancer Research : CR 41: 28. doi:10.1186/s13046-021-02211-2.

56. Martina, José A., Heba I. Diab, Owen A. Brady, and Rosa Puertollano. 2016. “TFEB and TFE3 Are Novel Components of the Integrated Stress Response.” The EMBO journal 35(5): 479–95. doi:10.15252/embj.201593428.

57. Martina, José A., Heba I. Diab, Li Lishu, Lim Jeong-A, Simona Patange, Nina Raben, and Rosa Puertollano. 2014. “The Nutrient-Responsive Transcription Factor TFE3, Promotes Autophagy, Lysosomal Biogenesis, and Clearance of Cellular Debris.” Science signaling 7(309): ra9. doi:10.1126/scisignal.2004754.

58. Mathur, Pallavi, Camilla De Barros Santos, Hugo Lachuer, Julie Patat, Bruno Latgé, François Radvanyi, Bruno Goud, and Kristine Schauer. 2023. “Transcription Factor EB Regulates Phosphatidylinositol-3-Phosphate Levels That Control Lysosome Positioning in the Bladder Cancer Model.” Communications Biology 6: 114. doi:10.1038/s42003-023-04501-1.

59. Matthew-Onabanjo, Asia N., Jenny Janusis, Jose Mercado-Matos, Anne E. Carlisle, Dohoon Kim, Fayola Levine, Peter Cruz-Gordillo, et al. 2020. “Beclin 1 Promotes Endosome Recruitment of Hepatocyte Growth Factor Tyrosine Kinase Substrate to Suppress Tumor Proliferation.” Cancer Research 80(2): 249–62. doi:10.1158/0008-5472.CAN-19-1555.

60. Menon, Dilip, Apoorva Bhapkar, Bhoomika Manchandia, Gitanjali Charak, Surabhi Rathore, Rakesh Mohan Jha, Arpita Nahak, et al. 2023. “ARL8B Mediates Lipid Droplet Contact and Delivery to Lysosomes for Lipid Remobilization.” Cell Reports 42(10): 113203. doi:10.1016/j.celrep.2023.113203.

61. Mikhajlov, Oleg, Ram M. Adar, Maria Tătulea-Codrean, Anne-Sophie Macé, John Manzi, Fanny Tabarin, Aude Battistella, et al. 2025. “Cell Adhesion and Spreading on Fluid Membranes through Microtubules-Dependent Mechanotransduction.” Nature Communications 16(1): 1201. doi:10.1038/s41467-025-56343-6.

62. Moreno-Layseca, Paulina, Jaroslav Icha, Hellyeh Hamidi, and Johanna Ivaska. 2019. “Integrin Trafficking in Cells and Tissues.” Nature cell biology 21(2): 122–32. doi:10.1038/s41556-018-0223-z.

63. Orlikowska-Rzeznik, Hanna, Emilia Krok, Madhurima Chattopadhyay, Agnieszka Lester, and Lukasz Piatkowski. 2023. “Laurdan Discerns Lipid Membrane Hydration and Cholesterol Content.” The Journal of Physical Chemistry. B 127(15): 3382–91. doi:10.1021/acs.jpcb.3c00654.

64. Ostrowski, Matias, Nuno B. Carmo, Sophie Krumeich, Isabelle Fanget, Graça Raposo, Ariel Savina, Catarina F. Moita, et al. 2010. “Rab27a and Rab27b Control Different Steps of the Exosome Secretion Pathway.” Nature Cell Biology 12(1): 19–30. doi:10.1038/ncb2000.

65. Otakhor, Kelly O., Mohd Ali Abbas Zaidi, Rebecca Oberley-Deegan, Moorthy Ponnusamy, and Micah B. Schott. 2025. “RAB5 NUCLEOTIDE BINDING PROMOTES β-OXIDATION TO FUEL HEPATOCELLULAR CARCINOMA CELL PROLIFERATION.” bioRxiv: The Preprint Server for Biology: 2025.08.20.670915. doi:10.1101/2025.08.20.670915.

66. Ouni, Emna, Alexis Peaucelle, Rasta Ghasemi, Francesco Facchinetti, Matthieu Opitz, Ludovic Bigot, Allan Sauvat, et al. 2025. “Mechanosensitive Interactions of Tumoroids with an Engineered Environment Promote Cell Proliferation and Enhance Drug Response Detection.” Cell Biomaterials 1(8). doi:10.1016/j.celbio.2025.100149.

67. Owen, Dylan M., Carles Rentero, Astrid Magenau, Ahmed Abu-Siniyeh, and Katharina Gaus. 2011. “Quantitative Imaging of Membrane Lipid Order in Cells and Organisms.” Nature Protocols 7(1): 24–35. doi:10.1038/nprot.2011.419.

68. Palmieri, Michela, Soren Impey, Hyojin Kang, Alberto di Ronza, Carl Pelz, Marco Sardiello, and Andrea Ballabio. 2011. “Characterization of the CLEAR Network Reveals an Integrated Control of Cellular Clearance Pathways.” Human molecular genetics 20(19): 3852–66. doi:10.1093/hmg/ddr306.

69. Parasassi, T., G. De Stasio, G. Ravagnan, R. M. Rusch, and E. Gratton. 1991. “Quantitation of Lipid Phases in Phospholipid Vesicles by the Generalized Polarization of Laurdan Fluorescence.” Biophysical Journal 60(1): 179–89. doi:10.1016/S0006-3495(91)82041-0.

70. Pastore, Nunzia, Owen A. Brady, Heba I. Diab, José A. Martina, Lu Sun, Tuong Huynh, Jeong-A Lim, et al. 2016. “TFEB and TFE3 Cooperate in the Regulation of the Innate Immune Response in Activated Macrophages.” Autophagy 12(8): 1240–58. doi:10.1080/15548627.2016.1179405.

71. Pastore, Nunzia, Anna Vainshtein, Tiemo J Klisch, Andrea Armani, Tuong Huynh, Niculin J Herz, Elena V Polishchuk, Marco Sandri, and Andrea Ballabio. 2017. “TFE3 Regulates Whole-body Energy Metabolism in Cooperation with TFEB.” EMBO Molecular Medicine 9(5): 605–21. doi:10.15252/emmm.201607204.

72. Peng, Wang, Shu Chen, Jingyu Ma, Wenjie Wei, Naixin Lin, Jinchao Xing, Wenjing Guo, et al. 2025. “Endosomal Trafficking Participates in Lipid Droplet Catabolism to Maintain Lipid Homeostasis.” Nature Communications 16(1): 1917. doi:10.1038/s41467-025-57038-8.

73. Perera, Rushika M., Chiara Di Malta, and Andrea Ballabio. 2019. “MiT/TFE Family of Transcription Factors, Lysosomes, and Cancer.” Annual review of cancer biology 3: 203–22. doi:10.1146/annurev-cancerbio-030518-055835.

74. Perera, Rushika M., Svetlana Stoykova, Brandon N. Nicolay, Kenneth N. Ross, Julien Fitamant, Myriam Boukhali, Justine Lengrand, et al. 2015. “Transcriptional Control of the Autophagy-Lysosome System in Pancreatic Cancer.” Nature 524(7565): 361–65. doi:10.1038/nature14587.

75. Pradat, Yoann, Julien Viot, Andrey A. Yurchenko, Konstantin Gunbin, Luigi Cerbone, Marc Deloger, Guillaume Grisay, et al. 2023. “Integrative Pan-Cancer Genomic and Transcriptomic Analyses of Refractory Metastatic Cancer.” Cancer Discovery 13(5): 1116–43. doi:10.1158/2159-8290.CD-22-0966.

76. Radulovic, Maja, Chonglin Yang, and Harald Stenmark. 2026. “Lysosomal Membrane Homeostasis and Its Importance in Physiology and Disease.” Nature Reviews. Molecular Cell Biology 27(1): 71–87. doi:10.1038/s41580-025-00873-w.

77. Raiborg, Camilla, Bjørn Bremnes, Anja Mehlum, David J. Gillooly, Antonello D’Arrigo, Espen Stang, and Harald Stenmark. 2001. “FYVE and Coiled-Coil Domains Determine the Specific Localisation of Hrs to Early Endosomes.” Journal of Cell Science 114(12): 2255–63. doi:10.1242/jcs.114.12.2255.

78. Raiborg, Camilla, Kristi Grønvold Bache, Anja Mehlum, Espen Stang, and Harald Stenmark. 2001. “Hrs Recruits Clathrin to Early Endosomes.” The EMBO Journal 20(17): 5008–21. doi:10.1093/emboj/20.17.5008.

79. Ruiz, Mario, Marcus Ståhlman, Jan Borén, and Marc Pilon. 2019. “AdipoR1 and AdipoR2 Maintain Membrane Fluidity in Most Human Cell Types and Independently of Adiponectin.” Journal of Lipid Research 60(5): 995–1004. doi:10.1194/jlr.M092494.

80. Salma, Nunciada, Jun S. Song, Zoltan Arany, and David E. Fisher. 2015. “Transcription Factor Tfe3 Directly Regulates Pgc-1alpha in Muscle.” Journal of Cellular Physiology 230(10): 2330–36. doi:10.1002/jcp.24978.

81. Sampaio, Julio L., Mathias J. Gerl, Christian Klose, Christer S. Ejsing, Hartmut Beug, Kai Simons, and Andrej Shevchenko. 2011. “Membrane Lipidome of an Epithelial Cell Line.” Proceedings of the National Academy of Sciences of the United States of America 108(5): 1903–7. doi:10.1073/pnas.1019267108.

82. Sardiello, Marco, Michela Palmieri, Alberto di Ronza, Diego Luis Medina, Marta Valenza, Vincenzo Alessandro Gennarino, Chiara Di Malta, et al. 2009. “A Gene Network Regulating Lysosomal Biogenesis and Function.” Science (New York, N.Y.) 325(5939): 473–77. doi:10.1126/science.1174447.

83. Sass, Frederike, Christian Schlein, Michelle Y. Jaeckstein, Paul Pertzborn, Michaela Schweizer, Thorsten Schinke, Andrea Ballabio, et al. 2021. “TFEB Deficiency Attenuates Mitochondrial Degradation upon Brown Adipose Tissue Whitening at Thermoneutrality.” Molecular Metabolism 47: 101173. doi:10.1016/j.molmet.2021.101173.

84. Scheinpflug, Kathi, Oxana Krylova, and Henrik Strahl. 2017. “Measurement of Cell Membrane Fluidity by Laurdan GP: Fluorescence Spectroscopy and Microscopy.” Methods in Molecular Biology 1520: 159–74. doi:10.1007/978-1-4939-6634-9_10.

85. Settembre, Carmine, Rossella De Cegli, Gelsomina Mansueto, Pradip K. Saha, Francesco Vetrini, Orane Visvikis, Tuong Huynh, et al. 2013. “TFEB Controls Cellular Lipid Metabolism through a Starvation-Induced Autoregulatory Loop.” Nature cell biology 15(6): 647–58. doi:10.1038/ncb2718.

86. Settembre, Carmine, Chiara Di Malta, Vinicia Assunta Polito, Moises Garcia Arencibia, Francesco Vetrini, Serkan Erdin, Serpil Uckac Erdin, et al. 2011. “TFEB Links Autophagy to Lysosomal Biogenesis.” Science (New York, N.Y.) 332(6036): 1429–33. doi:10.1126/science.1204592.

87. Sezgin, Erdinc, Ilya Levental, Satyajit Mayor, and Christian Eggeling. 2017. “The Mystery of Membrane Organization: Composition, Regulation and Roles of Lipid Rafts.” Nature Reviews. Molecular Cell Biology 18(6): 361–74. doi:10.1038/nrm.2017.16.

88. Shafaq-Zadah, Massiullah, Carina S. Gomes-Santos, Sabine Bardin, Paolo Maiuri, Mathieu Maurin, Julian Iranzo, Alexis Gautreau, et al. 2016. “Persistent Cell Migration and Adhesion Rely on Retrograde Transport of β(1) Integrin.” Nature Cell Biology 18(1): 54–64. doi:10.1038/ncb3287.

89. Song, Pei Xuan, Juan Peng, Mohyeddine Omrane, Ting Ting Cai, Didier Samuel, and Ama Gassama-Diagne. 2022. “Septin 9 and Phosphoinositides Regulate Lysosome Localization and Their Association with Lipid Droplets.” iScience 25(5): 104288. doi:10.1016/j.isci.2022.104288.

90. Taelman, Vincent F., Radoslaw Dobrowolski, Jean-Louis Plouhinec, Luis C. Fuentealba, Peggy P. Vorwald, Iwona Gumper, David D. Sabatini, and Edward M. De Robertis. 2010. “Wnt Signaling Requires the Sequestration of Glycogen Synthase Kinase 3 inside Multivesicular Endosomes.” Cell 143(7): 1136–48. doi:10.1016/j.cell.2010.11.034.

91. Thelen, Ashley M., and Roberto Zoncu. 2017. “Emerging Roles for the Lysosome in Lipid Metabolism.” Trends in Cell Biology 27(11): 833–50. doi:10.1016/j.tcb.2017.07.006.

92. Veatch, Sarah L., and Sarah L. Keller. 2002. “Organization in Lipid Membranes Containing Cholesterol.” Physical Review Letters 89(26): 268101. doi:10.1103/PhysRevLett.89.268101.

93. Wang, Yanning, Shulin Li, Marcel Mokbel, Alexander I. May, Zizhen Liang, Yonglun Zeng, Weiqi Wang, et al. 2024. “Biomolecular Condensates Mediate Bending and Scission of Endosome Membranes.” Nature 634(8036): 1204–10. doi:10.1038/s41586-024-07990-0.

94. Wang, Yun-Ting, Jiajie Chen, Xiang Li, Michihisa Umetani, Yang Chen, Pin-Lan Li, and Yang Zhang. 2019. “Contribution of Transcription Factor EB to adipoRon-Induced Inhibition of Arterial Smooth Muscle Cell Proliferation and Migration.” American Journal of Physiology. Cell Physiology 317(5): C1034–47. doi:10.1152/ajpcell.00294.2019.

95. Wei, Shuanzeng, Joseph R. Testa, and Pedram Argani. 2022. “A Review of Neoplasms with MITF/MiT Family Translocations.” Histology and Histopathology 37(4): 311–21. doi:10.14670/HH-18-426.

96. Yu, Kaikai, Guan M. Wang, Shiny Shengzhen Guo, Florian Bassermann, and Reinhard Fässler.2024. “The USP12/46 Deubiquitinases Protect Integrins from ESCRT-Mediated Lysosomal Degradation.” EMBO reports 25(12): 5687–5718. doi:10.1038/s44319-024-00300-9.

97. Zhao, G Q, Q Zhao, X Zhou, M G Mattei, and B de Crombrugghe. 1993. “TFEC, a Basic Helix-Loop-Helix Protein, Forms Heterodimers with TFE3 and Inhibits TFE3-Dependent Transcription Activation.” Molecular and Cellular Biology 13(8): 4505–12. doi:10.1128/mcb.13.8.4505.

98. Zhu, Xuejin, Yangjia Zhuo, Shulin Wu, Yanfei Chen, Jianheng Ye, Yulin Deng, Yuanfa Feng, et al. 2021. “TFEB Promotes Prostate Cancer Progression via Regulating ABCA2-Dependent Lysosomal Biogenesis.” Frontiers in Oncology 11. doi:10.3389/fonc.2021.632524.

99. Zoncu, Roberto, and Rushika M. Perera. 2023. “Emerging Roles of the MiT/TFE Factors in Cancer.” Trends in Cancer 9(10): 817–27. doi:10.1016/j.trecan.2023.06.005.

